# PP2A methylesterase, PME-1, and PP2A methyltransferase, LCMT-1, control sensitivity to impairments caused by injury-related oligomeric tau

**DOI:** 10.1101/2025.06.05.657660

**Authors:** Sowmya N. Sundaresh, Edward W. Vogel, Christopher D. Hue, Hong Zhang, Anna Staniszewski, Hanna L. Berman, Zafar Gill, Kesava Asam, Siqi Liang, Liwei Shen, Madhumathi Gnanaprakash, Erica Acquarone, Mauro Fà, Nicholas M. Kanaan, Barclay Morrison, Ottavio Arancio, Russell E. Nicholls

**Affiliations:** Department of Pathology and Cell Pathology, Columbia University, New York, NY, USA; Taub Institute for Research on Alzheimer’s Disease and the Aging Brain, Columbia University, New York, NY, USA; Department of Biomedical Engineering, Columbia University, New York, NY, USA; Department of Biology, Pace University, New York, NY, USA; Department of Translational Neuroscience, Michigan State University, Grand Rapids, MI, USA; Department of Medicine, Columbia University, New York, NY, USA

## Abstract

Oligomeric species of tau are a hallmark of multiple neurodegenerative diseases such as Alzheimer’s disease (AD) and chronic traumatic encephalopathy (CTE). Given the evidence implicating protein phosphatase 2A (PP2A) in the molecular pathogenesis of tau-related neurodegenerative disorders, we sought to determine whether manipulating the expression of enzymes that regulate PP2A activity, such as leucine carboxyl methyltransferase 1 (LCMT-1) and protein methyl esterase 1 (PME-1), might impact pathological responses to oligomeric tau. Here, we tested the effect of transgenic overexpression of LCMT-1 or PME-1 on cognitive and electrophysiological impairments caused by exposure to either recombinant oligomeric human tau or oligomeric tau prepared from mice subjected to blast-induced traumatic brain injury. We found that overexpression of LCMT-1 reduced sensitivity to tau-induced impairments, while overexpression of PME-1 increased sensitivity to these impairments. Moreover, we found that shockwave exposure increased the propensity of endogenous tau to form toxic oligomers. These results suggest that manipulating LCMT-1 or PME-1 activity may represent novel therapeutic approaches for disorders involving exposure to pathogenic forms of oligomeric tau.

## INTRODUCTION

Recent evidence suggests that soluble extracellular aggregates of tau play a role in the molecular pathogenesis of tauopathies, including Alzheimer’s disease (AD) and chronic traumatic encephalopathy (CTE). Tau is released from neurons in an activity dependent manner ^1-4^, and elevated levels of extracellular tau are detected in the cerebral spinal fluid of AD patients ^5,6^. In addition, elevated levels of interstitial tau are observed in the brains of traumatic brain injury (TBI) patients where they are predictive of adverse clinical outcomes ^7^. Extracellular application of tau preparations to cell and animal models elicits disease-related molecular, electrophysiological, and cognitive changes in these systems ^3,8-14^, and treatment with antibodies to extracellular tau ameliorates tau-related impairments in animal tauopathy models ^15^.

Multiple lines of evidence suggest that the serine/threonine phosphatase, PP2A, plays a key role in the molecular pathogenesis of tauopathies. PP2A expression and activity were reduced in the post-mortem brains of AD patients ^16-19^, and genetically or pharmacologically reducing PP2A activity alone is sufficient to elicit tau histopathology and cognitive deficits in animal models ^20-26^. In addition, PP2A dysregulation resulting from impaired methylation of the PP2A catalytic subunit is one of the potential molecular mechanisms contributing to increased AD risk in hyperhomocysteinemic patients ^27^. Finally, pharmacological activation of PP2A reduces cognitive impairment and pathology in rodent AD and tauopathy models, as well as in in animal models of TBI ^28-33^.

One of the mechanisms by which PP2A is thought to contribute to tauopathies is through its role as the principal phosphatase for pathologically phosphorylated forms of tau ^34,35^. This diversity of PP2A substrates is mirrored by the large number of PP2A isoforms. Mature PP2A holoenzymes are composed of 3 different subunits, each encoded by multiple genes with different isoforms that are together capable of generating over 100 different possible subunit combinations ^36^. The regulation of PP2A is complex, involving posttranslational modifications, endogenous protein inhibitors, and regulated assembly and disassembly of its constituent subunits ^36-40^. One of the mechanisms by which PP2A is regulated is through methylation of the C-terminal leucine residue of its catalytic subunit. This methylation is controlled through the actions of leucine carboxyl methyltransferase 1, LCMT-1 (addition of methylation), and protein methyl esterase 1, PME-1 (removal of methylation) ^41-46^.

Previously, we found that altering the expression of LCMT-1 or PME-1 altered the pathogenic response to oligomeric amyloid beta ^47-49^, another hallmark of AD. Since extracellularly applied soluble oligomeric tau elicits similar disease-related cognitive and electrophysiological impairments ^3,8^, we sought to determine whether transgenic overexpression of LCMT-1 or PME-1 might also alter the pathogenic response to soluble oligomeric tau. In these experiments, we used recombinant oligomeric tau preparations as well as oligomeric tau prepared from shockwave-exposed mice. In both cases, we found that transgenic LCMT-1 overexpression protected against the cognitive and electrophysiological impairments caused by oligomeric tau exposure, while transgenic PME-1 overexpression increased sensitivity to these impairments. In the case of tau prepared from shockwave-exposed mice, we also found that tau pathogenicity was dependent on tau oligomer formation, and that shockwave exposure increased the propensity of tau to form these oligomeric species. These data suggest that manipulating LCMT-1 or PME-1 activity may represent novel viable therapeutic approaches to protect against the pathogenic effects of extracellular tau oligomers.

## RESULTS

### LCMT-1 overexpression protects mice from cognitive and electrophysiological impairments caused by exposure to recombinant oligomeric tau

We found previously that acute infusion of oligomeric recombinant human 4R/2N tau into the brains of mice elicited impairments in short-term spatial memory tasks that are used to model the types of hippocampus-dependent cognitive impairments exhibited by patients with AD^3^. To determine whether LCMT-1 might regulate the sensitivity of mice to these types of tau-induced impairments, we infused LCMT-1 overexpressing transgenic mice ^48^ along with their control siblings with oligomeric recombinant tau or vehicle and tested their performance on a 2-day radial arm water maze task (RAWM) ^50^. In this task, animals navigate to an escape platform in a fixed location at the end of one of the maze arms, and the number of errors or wrong-arm entries committed during each block of training trials is used as a measure of their ability to learn and remember the platform location. While administration of recombinant oligomeric tau significantly impaired the performance of control animals, the performance of the tau-infused LCMT-1 overexpressing animals was comparable to that of vehicle treated animals, suggesting that LCMT-1 overexpression protects against cognitive impairments caused by extracellularly applied tau (Fig. 1A). The performance of the tau-infused LCMT-1 overexpressing animals in this task did not appear to result from a general enhancement in cognitive performance, since vehicle-treated LCMT-1 overexpressing animals performed comparably to vehicle-treated controls in this task (Fig. 1A). Visible platform water maze task was used to test for potential differences in visual perception, motivation, or swimming ability among these groups. There were no significant differences in either escape latency (Fig. S1A), or swimming speed (Fig. S1B), suggesting that differences in these variables do not account for the differences observed among groups in the 2-day RAWM task.

**Fig. 1.**
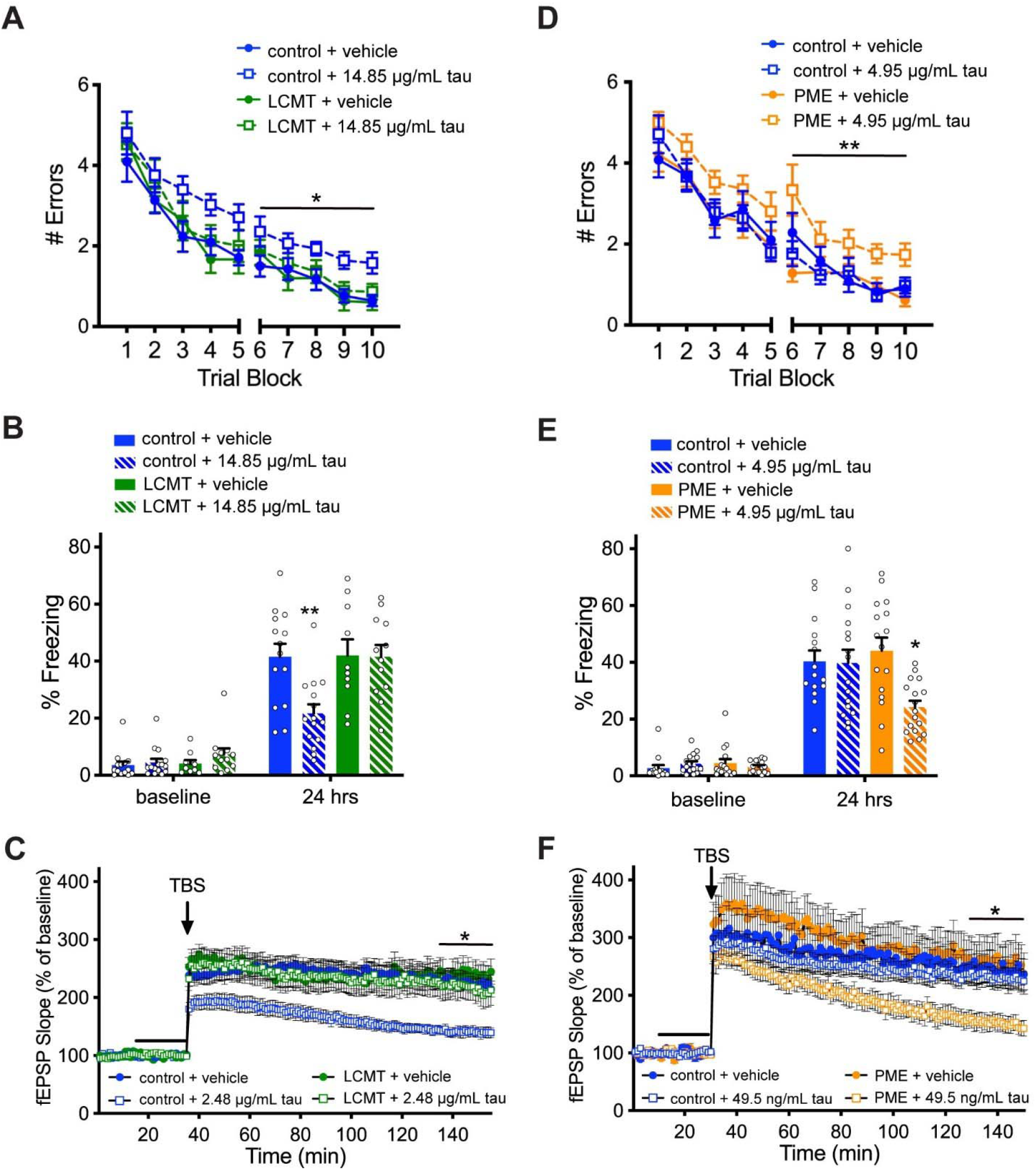
LCMT overexpression protects against, while PME overexpression increases sensitivity to behavioral and electrophysiological impairments caused by recombinant oligomeric tau. A) Number of errors committed during each 3-trial training block of a 2-day RAWM task for LCMT-1 overexpressing transgenic mice and sibling controls infused with recombinant oligomeric tau or vehicle. 2-way RM ANOVA for errors on day 2 (blocks 6-10) with group and block as factors shows a significant effect of group (F(3,47) = 6.995, P = 0.0006). Dunnett’s post-hoc comparisons show that tau-treated controls were significantly different from each of the other groups (P = 0.0006 vs. control+vehicle, P = 0.0017 vs. LCMT+vehicle, P = 0.0190 vs. LCMT+tau). N = 14 control+vehicle, 15 control+tau, 10 LCMT+vehicle, 12 LCMT+tau mice for this and the following panel. **B)** Percent of time spent freezing during initial exposure to the training context (baseline) and 24 hrs after foot shock for LCMT-1 overexpressing transgenic mice and sibling controls infused with recombinant oligomeric tau or vehicle. ANOVA for freezing at 24 hrs showed a significant difference among groups (F(3,47) = 6.015, P = 0.0015). Dunnett’s post-hoc comparisons show that tau-treated controls were significantly different from each of the other groups (P = 0.0030 vs. control+vehicle, P = 0.0061 vs. LCMT+vehicle, P = 0.0046 vs. LCMT+tau). No differences in baseline freezing were observed among groups on day 1 (ANOVA: F(3,47) = 1.244, P = 0.3045). **C)** Time course of Schaffer collateral fEPSP responses in hippocampal slices prepared from LCMT1 overexpressing transgenic mice and sibling controls and treated with recombinant oligomeric tau or vehicle for 20 min (black bar) prior to delivery of theta-burst stimulation (arrow). 2 -way RM ANOVA for fEPSP responses over the last 20 min of recording with group and block as factors shows a significant effect of group (F(3,47) = 4.112, P = 0.0114). Dunnett’s post-hoc comparisons show that tau-treated controls were significantly different from each of the other groups (P = 0.0098 vs. control+vehicle, P = 0.0116 vs. LCMT+vehicle, P = 0.0368 vs. LCMT+tau). N = 18 control+vehicle, 11 control+tau, 9 LCMT+vehicle, 13 LCMT+tau slices. **D)** Number of errors committed during each 3-trial training block of a 2-day RAWM task for PME-1 overexpressing transgenic mice and sibling controls infused with subthreshold doses of recombinant oligomeric tau or vehicle. 2-way RM ANOVA for errors on day 2 (blocks 6-10) with group and block as factors shows a significant effect of group (F(3,60) = 5.738, P = 0.0016). Dunnett’s post-hoc comparisons show that tau-treated PME overexpressing mice were significantly different from each of the other groups (P = 0.0058 vs. control+vehicle, P = 0.0125 vs control+tau, P = 0.0013 vs. PME+vehicle). N = 15 control+vehicle, 16 control+tau, 16 PME+vehicle, 17 PME+tau mice for this and the following panel. **E)** Percent of time spent freezing during initial exposure to the training context (baseline) and 24 hrs after foot shock for PME-1 overexpressing transgenic mice and sibling controls infused with subthreshold doses of recombinant oligomeric tau or vehicle. ANOVA for freezing at 24 hrs showed a significant difference among groups (F(3,60) = 5.255, P = 0.0028). Dunnett’s post-hoc comparisons show that tau-treated PME overexpressing mice were significantly different from each of the other groups (P = 0.0141 vs. control+vehicle, P = 0.0167 vs. control+tau, P = 0.0016 vs. PME+vehicle). No differences in baseline freezing were observed among groups on day 1 (ANOVA: F(3,60) = 0.6862, P = 0.5640). **F)** Time course of Schaffer collateral fEPSP responses in hippocampal slices prepared from PME-1 overexpressing transgenic mice and sibling controls and treated with subthreshold doses of recombinant oligomeric tau or vehicle for 20 min (black bar) prior to delivery of theta-burst stimulation (arrow). 2-way RM ANOVA for fEPSP responses over the last 20 min of recording with group and block as factors shows a significant effect of group (F(3,46) = 4.174, P = 0.0107). Dunnett’s post-hoc comparisons show that tau-treated PME overexpressing slices were significantly different from each of the other groups (P = 0.0213 vs. control+vehicle, P = 0.0304 vs. control+tau, P = 0.0100 vs. PME+vehicle). N = 11 control+vehicle, 16 control+tau, 10 PME+vehicle, 13 PME+tau slices. All data presented as mean ± SEM.

A contextual fear conditioning task was used as an additional test of the ability of LCMT-1 overexpression to protect against tau-induced cognitive impairments. In this task, animals learn to make a hippocampus-dependent association between a novel context and an aversive foot shock ^51^. As was the case for the 2-day RAWM task, we found that LMCT-1 overexpression protected mice from impairments in contextual fear conditioning caused by tau infusion. Tau infused control animals showed significantly reduced freezing upon reintroduction at 24 hrs, while the freezing response of LCMT-1 overexpressing animals was comparable at this time to vehicle treated controls (Fig. 1B). No differences were observed in the baseline or conditioned freezing responses between vehicle treated LCMT-1 and control mice, suggesting LCMT-1 overexpression itself did not affect the freezing response (Fig. 1B). To test for possible differences in shock perception among these groups, we examined their responses to a range of shock intensities and found comparable thresholds for the first visible, gross motor, and vocal response to foot shocks among these groups (Fig. S1C). An open field task was used to test for possible differences in baseline activity levels or anxiety among these groups that might confound our interpretation of their behavior in the fear conditioning task. We found no significant differences in anxiety (time spent in the center, Fig. S1D) or ambulatory activity (total distance travelled, Fig. S1E) in the open field environment.

Activity-dependent changes in the efficacy of synaptic transmission are thought to underlie particular forms of learning and memory, and disruptions in the ability to effect these changes are thought to contribute to cognitive impairments in patients with AD and other neurodegenerative disorders ^52-54^. Since our behavioral data suggest that LCMT-1 overexpression protects against tau-induced cognitive impairments, we sought to determine whether LCMT-1 overexpression might also protect against the tau-induced impairments in synaptic plasticity. To do this, we performed extracellular field potential recordings of long-term potentiation (LTP) at Schaffer collateral synapses in acute hippocampal slice preparations from LCMT-1 overexpressing and control animals. As reported previously^3^, bath application of control slices with recombinant oligomeric tau for 20 min prior to administration of a theta-burst stimulus train (TBS) significantly impaired potentiated responses relative to vehicle-treated control slices. However, similarly treated slices from LCMT-1 overexpressing mice showed no such impairment (Fig. 1C). LCMT-1 overexpression did not appear to affect LTP in the absence of tau, since potentiated responses in slices from vehicle-treated LCMT-1 overexpressing mice were comparable to those in vehicle-treated control slices (Fig. 1C). There was no significant difference in the stimulus-response relationship between LCMT-1 overexpressing and control mice, suggesting LCMT-1 overexpression did not affect the response to stimulation (Fig. S1F).

### PME-1 overexpression sensitizes mice to cognitive and electrophysiological impairments caused by exposure to recombinant oligomeric tau

Our earlier studies demonstrated opposing effects of LCMT-1 and PME-1 overexpression on Aβ sensitivity that was consistent with the complimentary biochemical activities of these enzymes ^47-49^. To determine whether PME-1 overexpression has opposing effects on tau-induced impairments compared to LCMT-1 overexpression, PME-1 overexpressing and control mice were treated with a subthreshold dose of recombinant tau that did not induce changes in cognitive behavior of control mice. PME-1 overexpressing mice treated with tau committed significantly more errors in 2-day RAWM than either tau infused control, vehicle infused control, or vehicle infused PME-1 overexpressing mice (Fig. 1D). The impaired performance of the PME-1 overexpressing animals did not appear to result from differences in visual perception, motivation or swimming ability since both escape latency (Fig. S1G) and swimming speed (Fig. S1H) were similar among all groups. In contextual fear conditioning, tau treated control mice showed freezing responses 24 hrs after training that were comparable to that of vehicle treated control animals, while the mean freezing response of the tau-infused PME-1 overexpressing animals was significantly reduced compared to the other groups (Fig. 1E). The baseline freezing levels prior to foot shock were comparable among groups, and they exhibited comparable thresholds for the first visible, gross motor, and vocal responses to foot shocks (Fig. S1I). Baseline anxiety (entries into the center, Fig. S1J) and ambulatory activity (total distance travelled, Fig. S1K) were also comparable among groups when tested in a novel open field environment. These results, together with the 2-day RAWM performance, suggest PME-1 overexpression increases sensitivity to tau-induced cognitive impairments.

To test the possibility that PME-1 overexpression also increases sensitivity to tau-induced impairments in synaptic plasticity, we treated acute hippocampal slices from these animals and their control siblings with vehicle or a subthreshold dose of recombinant oligomeric tau for 20 min prior to theta-burst stimulation. Potentiated responses in control slices treated with tau at this concentration were comparable to vehicle-treated controls, however, treatment of PME-1 overexpressing slices at this concentration resulted in significantly reduced potentiation relative to both vehicle-treated PME-1 overexpressing slices and controls (Fig. 1F). A comparison of the slope of evoked responses at increasing stimulus intensities obtained in slices from PME-1 overexpressing and control mice revealed no significant differences in the stimulus-response relationship between these groups (Fig. S1L) that might confound interpretation of these results. Together, these data suggest that PME-1 overexpression increases sensitivity to tau-induced LTP impairments that may underlie the increased sensitivity of these animals to tau-induced cognitive impairments.

### Murine model of shockwave induced TBI

Pathogenic forms of tau are thought to play a role in cognitive impairments and neurodegeneration that can result from TBI. TBI leads to a number of disease-linked biochemical modifications to tau including increased phosphorylation, prolyl isomerization, and soluble oligomer formation ^55^; and multiple studies reported that tau isolated from injured brains can elicit cellular, behavioral, and electrophysiological impairments when applied to healthy cell and animal models ^12,56-58^. A role for tau in TBI-related impairments is also supported by the observation that the absence of tau in tau knockout mice confers some protection against TBI related cognitive impairments ^59^, as well as the observation that treatment with anti-tau antibodies similarly protects mice from TBI related impairments ^56,60^.

Given the role of tau in TBI, we wanted to determine what effect LCMT-1 and PME-1 have on TBI-related tau impairments. To do this, wildtype mice were exposed to a shockwave injury using a custom designed shock tube (Fig. 2A) as previously described ^61-63^. Pressure transducers located at the exit of the shock tube and within the animal holder confirmed that the animal’s body was shielded effectively (Fig. 2B). Shockwave exposure resulted in a significant increase in righting time compared to sham exposure in wildtype mice (Fig. 2C).

**Fig. 2.**
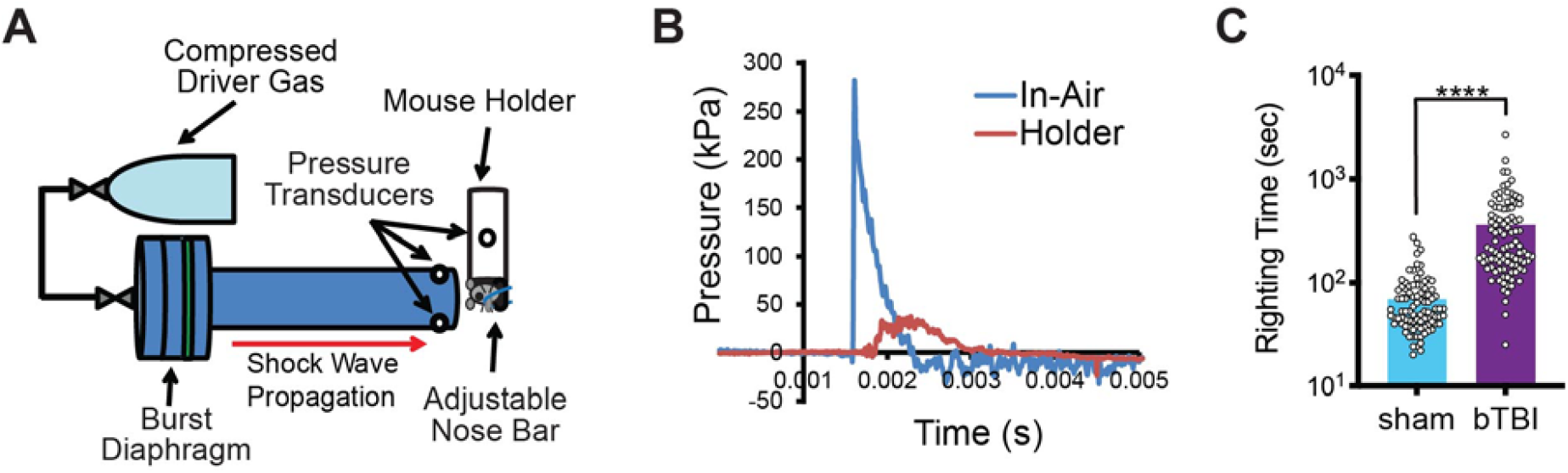
Murine shockwave exposure results in significantly increased righting time. **A)** Schematic diagram of shock tube setup and mouse holder. Device consisted of a cylinder of compressed helium connected to a driver section separated from a 1.2 m long, 76 mm diameter aluminum driven section by polyethylene terephthalate membranes (green). Animals were held in a brass/plexiglass tube lined with sorbothane that shielded the body, but left the head exposed. The head was positioned at the center of the tube approximately 1.5 cm from the exit with the dorsal surface perpendicular to the tube axis. The head was supported on the ventral surface by a sorbothane covered extension of the brass animal holder, and held in place with a wire nose bar. Two sensors located at the tube exit and one on the interior of the mouse holder recorded the pressure traces generated by the shockwave. **B)** Sample pressure trace generated over time by the shock wave at the tube exit (blue) and inside the animal holder (red). **C)** Righting time for shockwave (bTBI) and sham exposed mice showed a significant effect of treatment (unpaired, one-tailed t test: t = 7.699, P < 0.0001) (N = 91 sham, 96 bTBI). Data presented as mean ± SEM.

### Shockwave exposure increases tau phosphorylation and produces a modest cognitive deficit at 2 weeks but not 3 months post-injury

To assess the effect of shockwave exposure on tau phosphorylation, we performed western blots on hippocampal homogenates from shockwave exposed mice (Fig. 3A-D) using antibodies against three different phospho-tau epitopes: the PHF1 antibody (phospho-Ser396/404), the CP13 antibody (phospho-Ser202), and the AT270 antibody (phospho-Thr181). We found that PHF1 immunoreactivity was significantly elevated in homogenates prepared from shockwave exposed mice at 1 hr and 24 hrs after injury relative to controls, but returned to control levels by 3 weeks after injury (Fig. 3A). CP13 immunoreactivity was significantly elevated in shockwave exposed animals compared to controls at 1 hr, 24 hrs, and 3 weeks after injury with the largest increase occurring at 1hr (Fig. 3B). Finally, AT270 immunoreactivity was also significantly elevated at 1 hr, but not at 24 hrs, or 3 weeks after injury (Fig. 3C). There was no significant effect of injury on total tau levels (Fig. 3D). Together these data suggest that shockwave exposure results in a transient increase in phosphorylation at least three different disease-related phospho-tau epitopes, and is consistent with previously published reports of other rodent models of TBI ^55^.

**Fig. 3.**
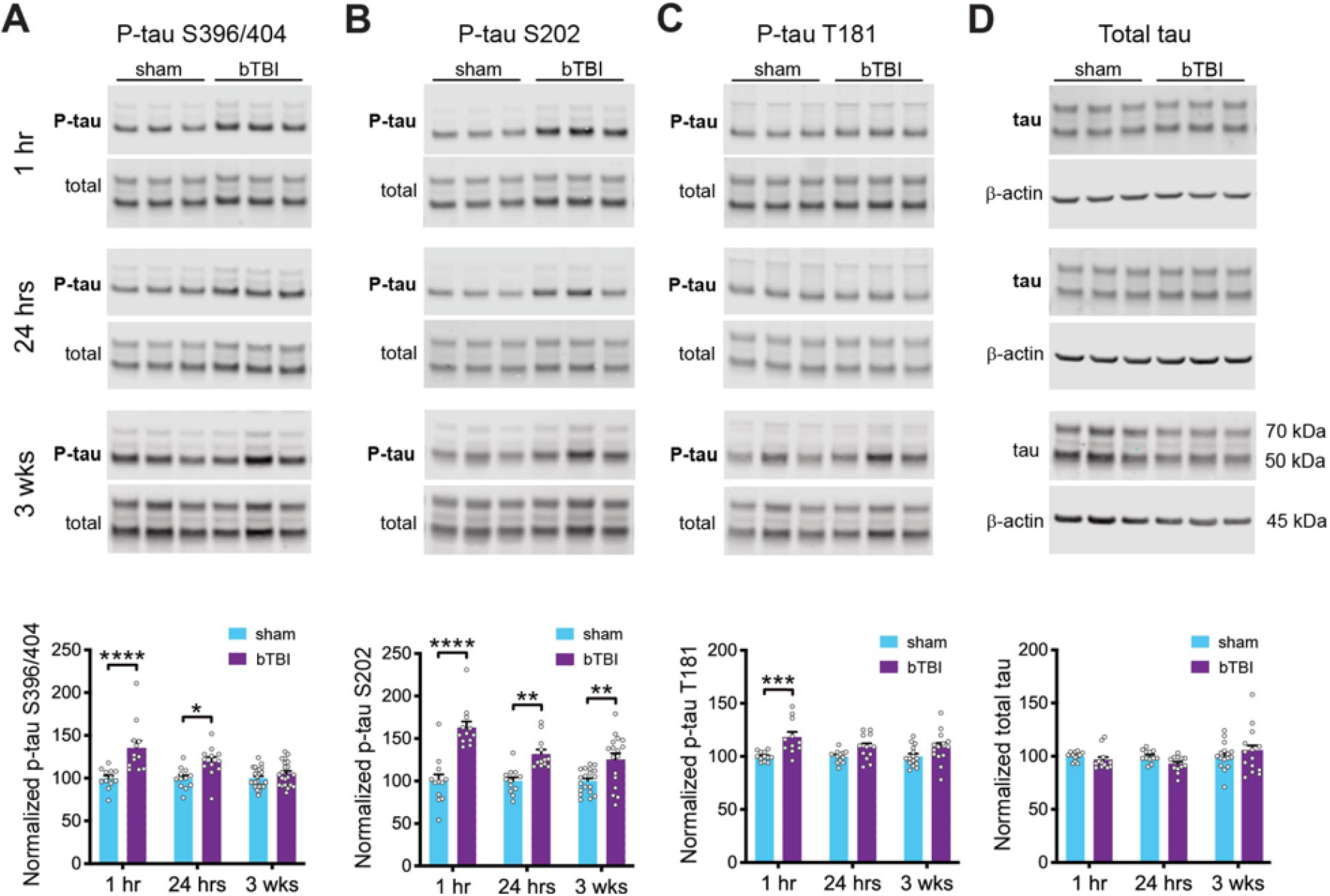
Shockwave exposure results in transient increase in tau phosphorylation. Representative western blots for phosphorylated tau (at serine 396/404, serine 202, and threonine 181), total tau and β-actin. **A)** Quantification of immunoreactivity for tau phosphorylated at serine 396/404 normalized to total tau in hippocampal homogenates from sham and shockwave (bTBI) exposed mice at 1 hr, 24 hrs, and 3 wks after injury, expressed as percent of control value. 2-way ANOVA with time point and treatment as factors shows a significant effect of treatment (F(1,86) = 31.29, P < 0.0001). Sidak’s post-hoc comparisons show significant differences between sham and bTBI groups at 1 hr (P < 0.0001) and 24 hrs (P = 0.0141), but not 3 wks (P = 0.6516). N=12 sham/12 bTBI for 1hr, 12 sham/12 bTBI for 24 hrs, and 23 sham/21 bTBI for 3 wks. **B)** Quantification of immunoreactivity for tau phosphorylated at serine 202 normalized to total tau in hippocampal homogenates from sham and shockwave exposed mice at 1 hr, 24 hrs, and 3 wks after injury, expressed as percent of control value. 2-way ANOVA with time point and treatment as factors shows a significant effect of treatment (F(1,78) = 64.04, P < 0.0001). Sidak’s post-hoc comparisons show significant differences between sham and bTBI groups at 1 hr (P < 0.0001), 24 hrs (P = 0.0025), and 3 wks (P = 0.0036). N = 12 sham/12 bTBI for 1hr, 12 sham/12 bTBI for 24 hrs, and 19 sham/17 bTBI for 3 wks. **C)** Quantification of immunoreactivity for tau phosphorylated at threonine 181 normalized to total tau in hippocampal homogenates from sham and shockwave exposed mice at 1 hr, 24 hrs, and 3 wks after injury, expressed as percent of control value. 2-way ANOVA with time point and treatment as factors shows a significant effect of treatment (F(1,70) = 20.00, P < 0.0001). Sidak’s post-hoc comparisons show significant differences between sham and bTBI groups at 1 hr (P = 0.0010), but not 24 hrs (P = 0.1738) or 3 wks (P = 0.1434). N = 12 sham/12 bTBI for 1hr, 12 sham/12 bTBI for 24 hrs, and 15 sham/14 bTBI for 3 wks. **D)** Quantification of immunoreactivity for total tau normalized to β-actin in hippocampal homogenates from sham and shockwave exposed mice at 1 hr, 24 hrs, and 3 wks after injury, expressed as percent of control value. 2-way ANOVA with time point and treatment as factors shows no significant effect of treatment (F(1,72) = 0.5707, P = 0.4524). Sidak’s post-hoc comparisons show significant differences between sham and bTBI groups at 1 hr (P = 0.8705), but not 24 hrs (P = 0.3940) or 3 wks (P = 0.7105). N = 12 sham/12 bTBI for 1hr, 12 sham/12 bTBI for 24 hrs, and 16 sham/14 bTBI for 3 wks. All data presented as mean ± SEM.

To further characterize this model, we assessed cognitive abilities of shockwave and sham exposed animals at multiple timepoints following injury, and found evidence that shockwave exposure under these conditions produces a modest, transient, cognitive impairment. In the contextual fear conditioning task, there were no significant differences in performance between the shockwave and sham-exposed animals at either 2 weeks or 3 months post-injury (Fig. S2 A,B). In the 2-day RAWM task, there was a significant increase in the number of errors committed by shockwave exposed mice at 2 weeks, but not 3 months after injury (Fig. S2E-F). This deficit was not accompanied by significant differences in either escape latency or swim speed when these animals were assessed in a visible platform water maze task (Fig. S2G-J), suggesting that the impairment we observed at 2 weeks post-injury in the 2-day RAWM task was the result of a transient impairment in short-term spatial memory similar to that reported by Beamer et al. ^63^.

Additional behavioral assessments of these shockwave exposed animals identified no significant injury related effects on motor function, anxiety, or depressive behavior. To assess the impact of our shockwave exposure protocol on motor function, we tested two separate groups of mice at 2 weeks and 3 months after injury on an accelerating rotarod task, and found no significant differences between shockwave and sham-exposed mice at either time point (Fig. S3A-C). To test for injury-related effects on anxiety related behavior, we tested mice at 2 weeks and 3 months after injury on both a novel open field environment and an elevated plus maze, and found no significant differences between groups on either apparatus at either time point (Fig. S3D-K). To test for injury-related increases in depressive behavior, we tested mice at 2 weeks and 3 months after injury in forced swim and tail suspension tasks, and similarly found no significant differences between groups in either task at either time point (Fig. S3L-O).

### Infusion of tau from shockwave-exposed mice elicits cognitive and electrophysiological impairments in wildtype mice

Published reports demonstrated that experimentally induced TBI leads to the production of pathogenic oligomeric tau species in rodent models, which can elicit cognitive and electrophysiological impairments in healthy animals and tissues ^55^. We sought to determine whether the same might be true for animals subjected to our shockwave-induced TBI protocol. To do this, we prepared tau from shockwave or sham-exposed brains as previously described ^3,64^. To control for the possibility that any activities we observed in the shockwave-exposed samples were due to an injury-related non-tau contaminant, we performed these same procedures in parallel on brain homogenates prepared from shockwave-exposed tau knockout (KO) animals.

We then infused tau from these preparations into the brains of naïve wild-type mice and tested their cognitive performance We found that animals infused with tau prepared from shockwave-exposed brains, referred to as blast tau, made significantly more errors in the 2-day RAWM task, and exhibited significantly lower freezing responses in the contextual fear conditioning task than vehicle-infused control animals (Fig. 4A,B). In contrast, the performance of animals infused with tau prepared from sham-exposed brains, referred to as sham tau, in these tasks was comparable to vehicle-infused animals. The performance of animals infused with material prepared from shockwave-exposed tau knockout brains, was also comparable to the vehicle-infused controls, suggesting that the ability of blast tau to impair cognitive performance was dependent on both shockwave exposure and tau. Moreover, the behavior impairments we observed in animals infused with blast tau was not accompanied by impairments in visible platform water maze performance, shock perception, or open field behavior, suggesting that they were not the result of changes in non-cognitive factors (Fig. S4A-E).

**Fig. 4.**
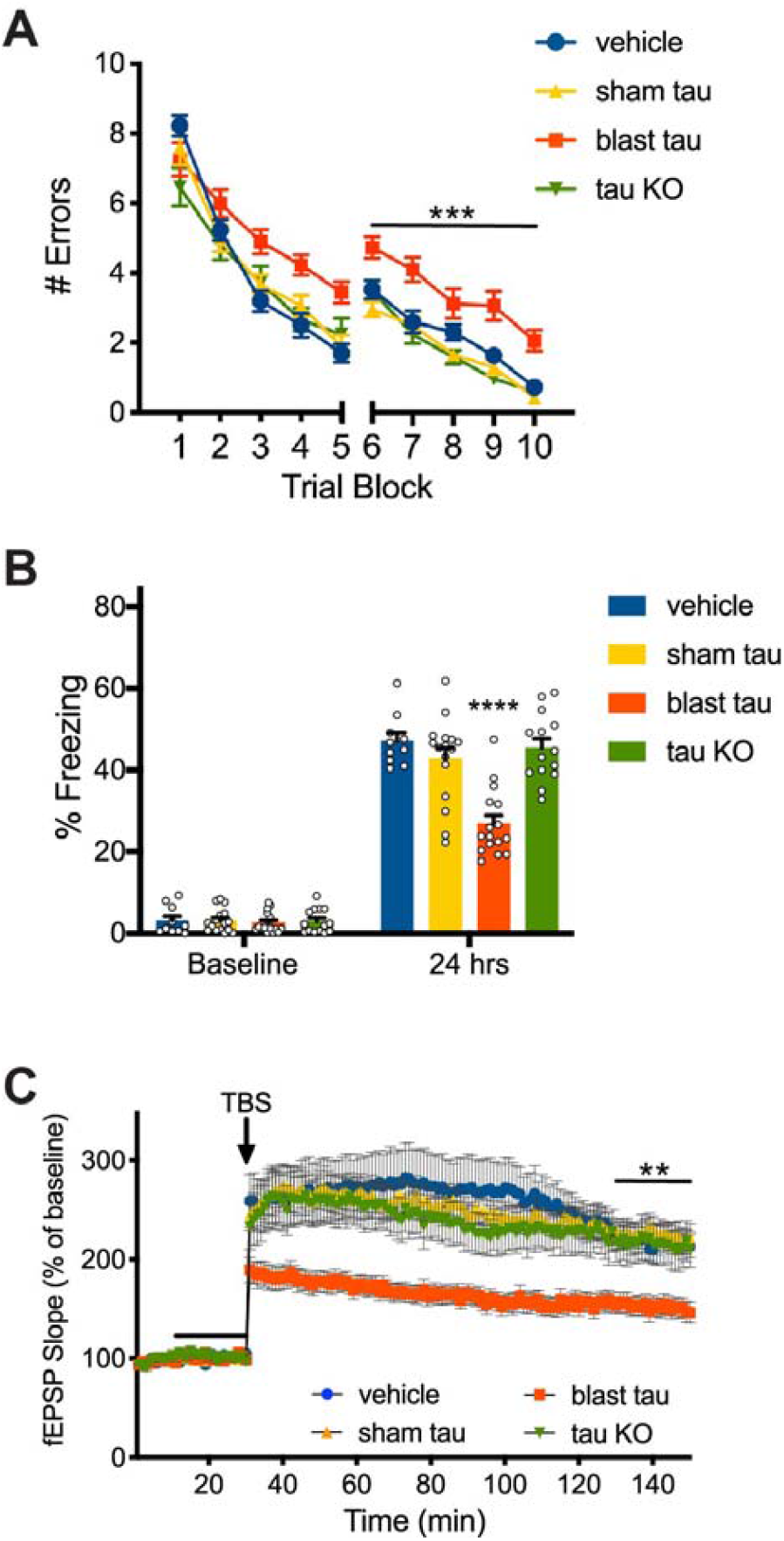
Tau from shockwave-exposed but not sham-exposed mice impairs cognition and synaptic plasticity. **A)** Number of errors committed during testing on a 2-day RAWM task for wild type mice infused with vehicle, tau isolated from sham-exposed mice (sham tau), tau isolated from shockwave-exposed mice (blast tau), or mock purified material from shockwave exposed tau KO mice (tau KO) shows that blast tau treated mice commit significantly more errors than each of the other groups. 2-way RM ANOVA for errors on day 2 (blocks 6-10) with group and block as factors shows a significant effect of group (F(3,52) = 16.41, P < 0.0001). Dunnett’s post-hoc comparisons show that blast tau treated mice were significantly different from vehicle-treated mice (P = 0.0005), sham tau-treated mice (P < 0.0001) and mock purified material from shockwave exposed tau KO mice (P < 0.0001). N = 10 vehicle, 16 sham tau, 16 blast tau, 14 tau KO treated mice for this and the following panel. **B)** Time spent freezing during initial pre-foot shock exposure to the fear conditioning chamber (baseline) and during reintroduction 24 hrs later for mice infused with vehicle, sham tau, blast tau, or mock purified material from shockwave exposed tau KO mice. No significant differences in baseline freezing were observed among groups (ANOVA: F(3,52) = 0.1207, P = 0.9475). However, one-way ANOVA for freezing at 24 hrs showed a significant difference among groups (F(3,52) = 16.90, P < 0.0001). Dunnett’s multiple comparisons show that the blast tau treated group was significantly different from each of the other three groups (P < 0.0001). **C)** Time course of Schaffer collateral fEPSP responses prior to and following delivery of theta-burst stimulation (arrow) in wild-type hippocampal slices treated with vehicle, sham tau, blast tau, or mock purified material from shockwave exposed tau KO mice for 20 min (black bar). 2-way RM ANOVA for fEPSP responses over the last 20 min of recording with treatment and time as factors shows a significant effect of treatment (F(3,54) = 8.796, P < 0.0001). Dunnett’s multiple comparisons show that the blast tau treated group was significantly different from each of the other three groups (vs. vehicle: P = 0.0039; vs. sham tau: P < 0.0001; vs. tau KO: P = 0.0064). N = 9 vehicle, 20 sham tau, 20 blast tau, 9 tau KO treated slices. All data presented as mean ± SEM.

To determine whether tau isolated from our shockwave-exposed animals might also interfere with activity-dependent changes in synaptic efficacy that underlie cognition, we treated acute hippocampal slices from wild-type mice with either vehicle, sham tau, blast tau, or preparation from shockwave-exposed tau KO mice for 20 min prior to theta-burst stimulation. We found that slices treated with blast tau significantly impaired LTP relative to vehicle-treated controls, while treatment with either sham tau or material from shockwave-exposed tau KO brains had no effect (Fig. 4C). As was the case for its ability to impair cognitive performance, the ability of the material isolated from shockwave-exposed brains to impair synaptic plasticity was, therefore, also dependent on both shockwave exposure and tau.

### Tau from shockwave-exposed mice impairs cognitive performance and synaptic plasticity in an oligomer-dependent manner

Data suggest that soluble tau oligomers contribute to impairments associated with TBI, and are responsible for the pathogenic activity observed in extracts prepared from injured brains ^55^. To determine whether the ability of our tau preparations to impair cognition and synaptic plasticity was dependent on tau’s oligomeric state, we used the TOC1 antibody that recognizes pathogenic oligomeric tau species ^65,66^. Normalization of TOC1 reactivity to total tau revealed a significant increase in oligomeric tau species in blast tau compared to sham tau preparations. (Fig. 5A). This is consistent with the hypothesis that oligomeric tau species are responsible for the cognitive and electrophysiological impairments produced by shockwave-exposed samples. Moreover, since sham tau did not lead to the production of TOC1 reactive tau species, these results also suggest that TBI under these conditions alters the propensity of tau to form these oligomeric species.

**Fig. 5.**
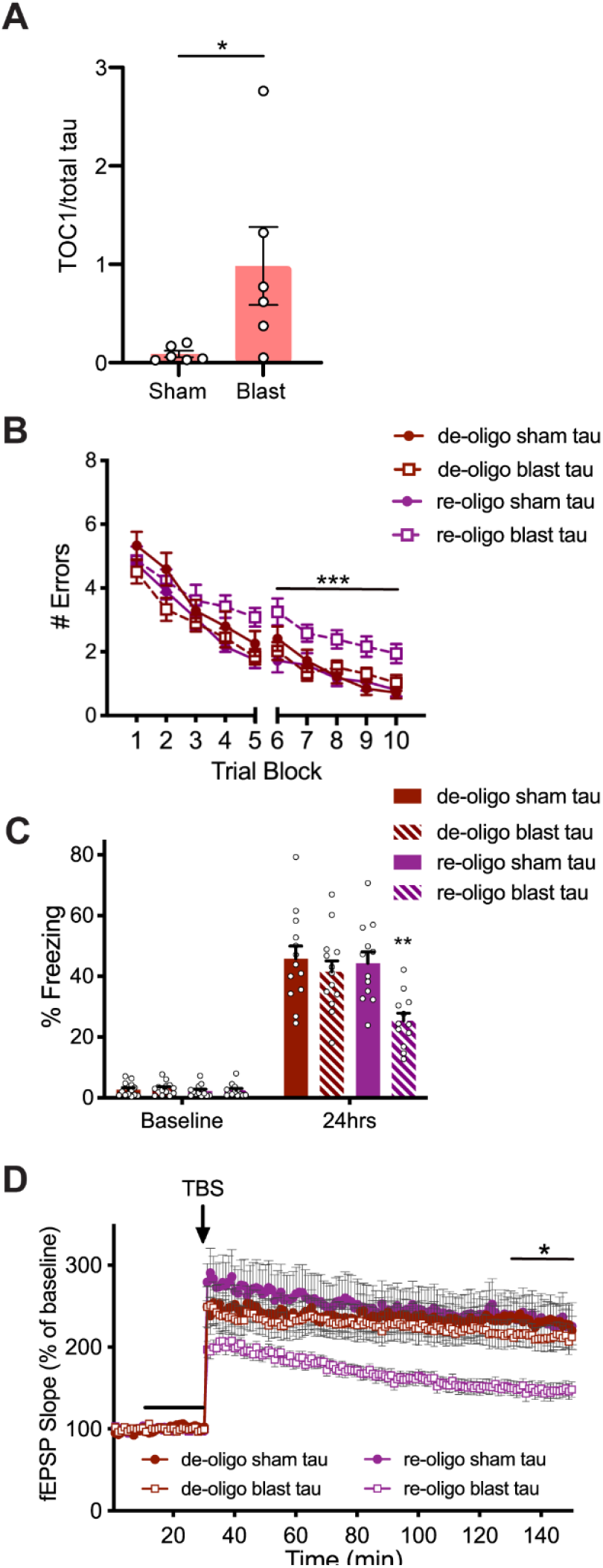
Blast tau impairs cognition and synaptic plasticity in an oligomerization-dependent manner. **A)** Graph representing TOC1 normalized to total tau based on sELISA shows a significant difference between tau isolated from shockwave-exposed mice (blast tau) and tau isolated from sham-exposed mice (sham tau) (unpaired, one-tailed t-test: t = 2.254, P = 0.0239). N = 6 sham tau, 6 blast tau. **B)** Number of errors committed during testing on a 2-day RAWM task for mice infused with sham or blast tau that was de-oligomerized by treatment with reducing reagent alone (de-oligo) or de-oligomerized then re-oligomerized by peroxide treatment (re-oligo). 2-way RM ANOVA for errors on day 2 (blocks 6-10) with group and block as factors shows a significant effect of group (F(3,46) = 10.47, P < 0.0001), and Dunnett’s post-hoc comparisons show that re-oligomerized blast tau treated mice were significantly different from each of the other groups (P = 0.0001 vs. de-oligo sham tau, P = 0.0003 vs. de-oligo blast tau, and P < 0.0001 vs. re-oligo sham tau). N = 13 de-oligo sham tau, 13 de-oligo blast tau, 12 re-oligo sham tau, 12 re-oligo blast tau treated mice for this and the following panel. **C)** Time spent freezing during initial pre-foot shock exposure to the fear conditioning chamber (baseline) and during reintroduction 24 hrs later for mice infused with de-oligo or re-oligo sham or blast tau. No significant differences in baseline freezing were observed among groups (ANOVA: F(3,46) = 0.3240, P = 0.8080). However, one-way ANOVA for freezing at 24 hrs showed a significant difference among groups (F(3,46) = 6.617, P = 0.0008). Dunnett’s post-hoc comparisons at 24 hrs show that re-oligomerized blast tau treated mice were significantly different from each of the other groups (P = 0.0007 vs. de-oligo sham tau, P = 0.0081 vs. de-oligo blast tau, and P = 0.0020 vs. re-oligo sham tau). **D)** Time course of Schaffer collateral fEPSP responses prior to and following delivery of theta-burst stimulation (arrow) in wild-type hippocampal slices treated with de-oligo or re-oligo sham or blast tau for 20 min (black bar). 2 -way RM ANOVA for fEPSP responses over the last 20 min of recording with treatment and time as factors shows a significant effect of treatment (F(3,62) = 5.371, P = 0.0024). Dunnett’s multiple comparisons show that the re-oligo blast tau treated group was significantly different from each of the other three groups (P = 0.0030 vs. de-oligo sham tau, P = 0.0250 vs. de-oligo blast tau, and P = 0.0040 vs. re-oligo sham tau). N = 17 de-oligo sham tau, 17 de-oligo blast tau, 14 re-oligo sham tau, 18 re-oligo blast tau treated slices. All data presented as mean ± SEM.

To verify the correlation between tau’s pathogenic activity and its oligomeric state further, we compared the effects of oligomerized and de-oligomerized tau preparations on cognitive performance and synaptic plasticity. Tau prepared from shockwave and sham-exposed brains were treated with dithiothreitol (DTT) and ethylenediaminetetraacetic acid (EDTA) to induce de-oligomerization and a fraction of this sample was treated with hydrogen peroxide again to induce re-oligomerization. Only infusion with re-oligomerized blast tau elicited impairments in the RAWM and contextual fear conditioning tasks (Fig. 5B,C). De-oligomerizing blast tau significantly reduced or eliminated its ability to impair the cognitive performance of wild-type mice, which was comparable to the performance of mice treated with sham tau regardless of whether the latter was de-oligomerized or re-oligomerized. None of these preparations had significant effects on behavior in the visible platform water maze task, sensory threshold assessment, or open field environment, suggesting the impairments produced by re-oligomerized blast tau in the RAWM and contextual fear conditioning tasks were not due to changes in non-cognitive factors that could have impacted performance in these tasks (Fig. S4F-J).

The effect of these preparations on LTP was similar to its effect on cognitive behavior (Fig. 5D). De-oligomerizing blast tau significantly reduced or eliminated its ability to impair LTP, while re-oligomerizing blast tau restored this ability. Consistent with their effects on cognitive performance, neither de-oligomerized, nor re-oligomerized sham tau elicited significant impairments of LTP, suggesting that the ability of blast tau to impair both cognition and synaptic plasticity are dependent on its oligomeric state.

### Overexpression of LCMT protects against while PME exacerbates oligomeric blast tau induced behavioral and electrophysiological impairments

We sought to determine whether the protective and sensitizing effects that LCMT-1 and PME-1 exhibit toward oligomeric recombinant tau extend to oligomeric tau derived from shockwave exposed mice. To assess the ability of LCMT-1 to protect against cognitive impairments produced by this form of oligomeric tau, we tested the behavior performance of LCMT-1 overexpressing mice and their control siblings that were treated with either sham or blast tau. We found that controls infused with blast tau made significantly more errors in the 2-day RAWM task, and exhibited significantly lower freezing responses in the contextual fear conditioning task than either the control or LCMT-1 overexpressing animals that were infused with sham tau (Fig. 6A,B). In contrast, the performance of LCMT-1 overexpressing animals that were infused with blast tau was comparable to the sham tau-treated groups in both of these tasks. To determine whether LCMT-1 overexpression might also protect against impairments in synaptic plasticity caused by exposure to blast tau, we compared the LTP response of acute hippocampal slices from LCMT-1 overexpressing and control mice treated with either sham or blast tau. We found that treatment of control slices with blast tau significantly impaired LTP relative to sham tau treated control and LCMT-1 overexpressing hippocampal slices, while LTP in the LCMT-1 overexpressing animals treated similarly with blast tau was comparable to sham tau treated groups (Fig. 6C).

**Fig. 6.**
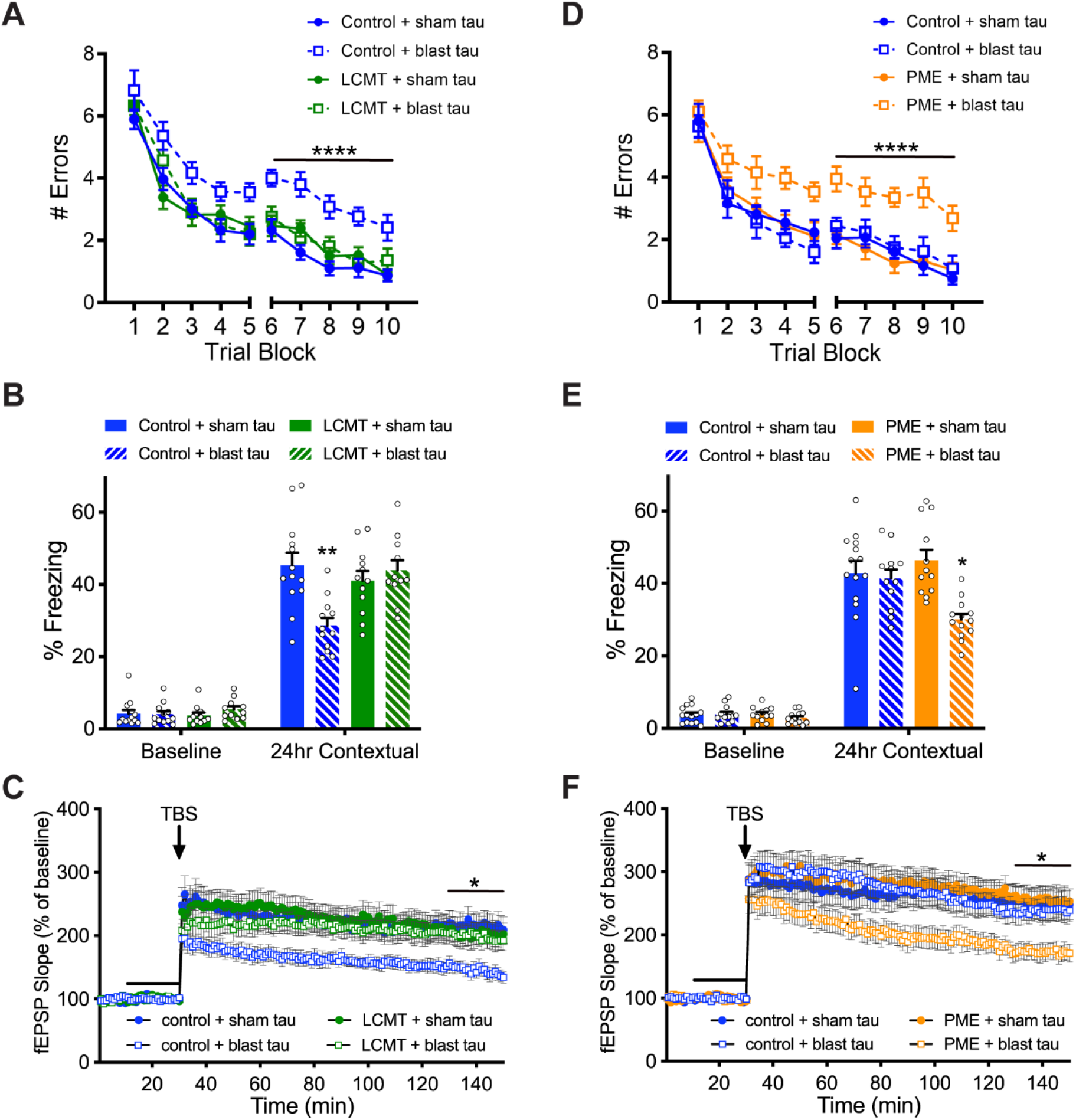
LCMT overexpression protects against, while PME overexpression increases sensitivity to behavioral and electrophysiological impairments caused by oligomeric tau prepared from shockwave-exposed mice. **A)** Number of errors committed during each 3-trial training block of a 2-day RAWM task for LCMT-1 overexpressing transgenic mice and sibling controls infused with sham or blast tau. 2-way RM ANOVA for errors on day 2 (blocks 6-10) with group and block as factors shows a significant effect of group (F(3,44) = 27.38, P < 0.0001). Dunnett’s post-hoc comparisons show that blast tau-treated controls were significantly different from each of the other groups (P < 0.0001). N = 13 control+sham tau, 12 control+blast tau, 12 LCMT+sham tau, 11 LCMT+blast tau mice for this and the following panel. **B)** Percent of time spent freezing during initial exposure to the training context (baseline) and 24 hrs after foot shock for LCMT-1 overexpressing transgenic mice and sibling controls infused with sham or blast tau. ANOVA for freezing at 24 hrs showed a significant difference among groups (F(3,44) = 7.245, P = 0.0005). Dunnett’s post-hoc comparisons show that blast tau-treated controls were significantly different from each of the other groups (P = 0.0003 vs. control+ sham tau, P = 0.0095 vs. LCMT+sham tau, P = 0.0016 vs. LCMT+blast tau). No differences in baseline freezing were observed among groups on day 1 (ANOVA: F(3,44) = 0.6219, P = 0.6046). **C)** Time course of Schaffer collateral fEPSP responses in hippocampal slices prepared from LCMT-1 overexpressing transgenic mice and sibling controls and treated with sham or blast tau for 20 min (black bar) prior to delivery of theta-burst stimulation (arrow). 2-way RM ANOVA for fEPSP responses over the last 20 min of recording with group and block as factors shows a significant effect of group (F(3,52) = 4.466, P = 0.0073). Dunnett’s post-hoc comparisons show that blast tau-treated controls were significantly different from each of the other groups (P = 0.0051 vs. control+sham tau, P = 0.0263 vs. LCMT+sham tau, P = 0.0333 vs. LCMT+blast tau). N = 13 control+sham tau, 16 control+blast tau, 13 LCMT+sham tau, 14 LCMT+blast tau slices. **D)** Number of errors committed during each 3-trial training block of a 2-day RAWM task for PME-1 overexpressing transgenic mice and sibling controls infused with subthreshold doses of sham or blast tau. Dunnett’s post-hoc comparisons show that blast tau-treated PME overexpressing mice were significantly different from each of the other groups (P < 0.0001). N = 14 control+sham tau, 12 control+blast tau, 12 PME+sham tau, 13 PME+blast tau mice for this and the following panel. **E)** Percent of time spent freezing during initial exposure to the training context (baseline) and 24 hrs after foot shock for PME-1 overexpressing transgenic mice and sibling controls infused with subthreshold doses of sham or blast tau. ANOVA for freezing at 24 hrs showed a significant difference among groups (F(3,47) = 6.934, P = 0.0006). Dunnett’s post-hoc comparisons show that blast tau-treated PME overexpressing mice were significantly different from each of the other groups (P = 0.0033 vs. control+sham tau, P = 0.0129 vs. control+blast tau, P = 0.0003 vs. PME+sham tau). No differences in baseline freezing were observed among groups on day 1 (ANOVA: F(3,47) = 0.6661, P = 0.5771). **F)** Time course of Schaffer collateral fEPSP responses in hippocampal slices prepared from PME-1 overexpressing transgenic mice and sibling controls and treated with subthreshold doses of sham or blast tau for 20 min (black bar) prior to delivery of theta-burst stimulation (arrow). 2-way RM ANOVA for fEPSP responses over the last 20 min of recording with group and block as factors shows a significant effect of group (F(3,55) = 4.470, P = 0.0070). Dunnett’s post-hoc comparisons show that blast tau-treated PME overexpressing slices were significantly different from each of the other groups (P = 0.0110 vs. control+sham tau, P = 0.0272 vs. control+blast tau, P = 0.0093 vs. PME+sham tau). N = 14 control+sham tau, 17 control+blast tau, 12 PME+sham tau, 16 PME+blast tau slices. All data presented as mean ± SEM.

To test whether PME-1 overexpression increased sensitivity to impairments produced by oligomeric tau derived from shockwave-exposed animals, we infused PME-1 overexpressing transgenic mice and their control siblings with a subthreshold dose of sham or blast tau that did not produce impairments in control mice. Treatment with blast tau sensitized PME-1 overexpressing mice to impairments in 2-day RAWM as well as contextual fear conditioning (Fig. 6D,E) and impaired LTP in hippocampal slices from PME-1 overexpressing mice (Fig. 6F).

The effects of LCMT-1 and PME-1 overexpression on cognitive impairments produced by oligomeric tau from shockwave exposed mice were not accompanied by impairments in visible platform water maze performance, shock perception, or open field behavior (Fig. S5). These results further support our observations from the recombinant tau experiments in that LCMT-1 overexpression protects against, while PME-1 overexpression increases sensitivity to cognitive and electrophysiological impairments caused by soluble oligomeric tau.

### Overexpression of PME or LCMT did not alter response to shockwave exposure

Given that LCMT-1 overexpression protected, and PME-1 overexpression sensitized mice to impairments caused by oligomeric tau from shockwave-exposed animals, we sought to determine whether overexpression of these transgenes might affect the response to shockwave exposure per se. However, measures in shockwave-exposed LCMT-1 and PME-1 overexpressing animals revealed no significant effects of these transgenes on injury related increases in tau phosphorylation (Fig. S6) and neither did behavioral measures in shockwave-exposed PME-1 overexpressing animals and controls reveal any injury related increases in behavioral or cognitive impairments (Fig. S7-8).

## DISCUSSION

Our study shows that transgenic overexpression of the PP2A methyl transferase, LCMT-1, protects mice from cognitive and electrophysiological impairments caused by exposure to soluble oligomers of the microtubule-associated protein, tau. We found that this was the case for tau oligomers prepared from recombinantly produced human 4R/2N tau and tau oligomers prepared from the brains of mice subjected to shockwave-induced TBI. Conversely, transgenic overexpression of the PP2A methyl esterase, PME-1, sensitized mice to cognitive and electrophysiological impairments caused by exposure to soluble oligomeric tau from these sources. The complementary effects of these transgenes are consistent with their complementary biochemical roles in controlling the protein methylation of PP2A ^41-46^. Given the evidence implicating soluble tau oligomers in the molecular pathogenesis of tauopathies, these data suggest that targeting LCMT-1 or PME-1 may be viable approaches for ameliorating the impairments caused by soluble tau oligomers in these disorders ^67-69^.

As outlined above, multiple lines of evidence support a role for PP2A serine/threonine phosphatases in tauopathies ^70-72^. Since LCMT-1 and PME-1 control PP2A C-terminal methylation, and thereby subunit composition, one obvious possibility for the mechanism of their effects on tau-induced impairments is through their regulation of PP2A activity ^41-46^. PP2A was identified as the principal phosphatase for pathologically phosphorylated forms of tau, suggesting a role for PP2A in the production of disease-related forms of tau. However, in our study, soluble oligomerized tau was applied exogenously. Therefore, if alterations in PP2A-dependent tau dephosphorylation do play a role in the protective and sensitizing effects of LCMT-1 and PME-1, then these enzymes must be altering tau phosphorylation in a way that affects the response to exogenously applied tau.

Alternatively, LCMT-1 and PME-1 overexpression may be acting through PP2A to alter the phosphorylation of one or more of the other substrates of this highly abundant phosphatase. Another particularly relevant candidate is the amyloid precursor protein, APP. PP2A regulates the phosphorylation of this protein at threonine 668, which has been implicated in AD ^78^. We previously showed that the level of phosphorylation at this site is altered in both LCMT-1 and PME-1 overexpressing mice ^48^. Moreover, we found that APP aids oligomeric tau internalization into neurons ^10^. Together, these observations raise the possibility that LCMT-1 and PME-1 overexpression may be regulating the response to extracellular tau oligomers by altering their APP-dependent internalization.

It is also possible that the protective actions of LCMT-1 and PME-1 are mediated through a target other than PP2A. PP4 and PP6 are two other serine/threonine phosphatases that are closely related to PP2A ^79^. While less is known about the disease related functions of PP4 and PP6, published data suggests the potential involvement of PP4 in neurodegenerative conditions via its interactions with tau, APP, and the PI3K, MAPK, Wnt and Toll-like receptor signaling pathways, and PP6 via its interaction with the interleukin-1 signaling pathway ^71^. Since LCMT-1 was reported to regulate both PP4 and PP6, and PME-1 was reported to regulate PP4 ^79-81^, it is possible that LCMT-1 and PME-1 dependent regulation of PP4 and/or PP6 contribute to their effects on susceptibility to tau-induced impairments.

Here, we tested the effect of LCMT-1 and PME-1 overexpression on impairments caused only by extracellularly applied tau. It remains unknown whether, and to what extent, these transgenes impact the effects of pathogenic forms of tau produced within cells, in different cell types, or in different brain regions. It will also be interesting to identify the relationship between the mechanism by which LCMT-1 and PME-1 alter sensitivity to tau-induced impairments, and the mechanisms by which tau contributes to normal physiological function.

Different tauopathies exhibit characteristic differences in the brain regions and cell types they affect, the tau isoforms involved, and the appearance of the insoluble tau aggregates they produce. Recently, differences among these disorders were found to extend to the level of the secondary structure of the tau proteins that are present in the filamentous tau aggregates ^82^. In our experiments, we tested soluble oligomeric tau aggregates produced from both recombinantly produced human 4R/2N tau, and endogenous tau from shockwave-exposed mice. While LCMT-1 and PME-1 overexpression altered sensitivity to impairments elicited by tau from both of these sources, it remains unknown whether the protective and sensitizing effects of these enzymes extend to other tau aggregate structures. Different tau aggregates also differ in their ability to seed different types of tau pathology in different brain regions, and it will be interesting to determine whether LCMT-1 and PME-1 overexpression affect tau-induced seeding and propagation ^82^.

Interestingly, we found that the shockwave exposure protocol we utilized impacted the propensity of tau to form pathogenic aggregates. Tau isolated from shockwave, but not sham-exposed mice produced TOC1 reactive tau aggregates *in vitro*, and the ability of tau from shockwave-exposed mice to elicit cognitive and electrophysiological impairments was dependent on its oligomeric state. Multiple post-translational modifications of tau are implicated in injury-related pathogenesis. Consistent with previous reports from both animal models and human patients, we observe increases in tau phosphorylation following TBI in our model. Characteristic increases in tau phosphorylation are also a hallmark of AD and other tauopathies, and increased phosphorylation at these sites was proposed to promote tau aggregation ^83,84^. Increases in cis prolyl isomerization of tau were documented following TBI in both humans and animal models ^56,60^. Reduced Pin1-dependent cis/trans prolyl isomerization of tau was implicated in AD ^85^. The involvement of the cis isomer of tau in TBI-related impairments is supported by the observation that treatment with cis tau antibody ameliorated injury induced cellular, histological, and behavioral changes in rodent models ^56,60^. The observation that cis tau is more resistant to dephosphorylation and more prone to aggregation, suggests a model in which injury related increases in cis tau isomerization lead to an increase pathologically phosphorylated cis tau that then aggregates to form pathogenic oligomers ^86^. However, the increased propensity of tau from shockwave exposed mice to form pathogenic aggregates may also be related to other post-translational modifications that have been described for tau ^87,88^.

Our data show that LCMT-1 and PME-1 regulate sensitivity to cognitive and electrophysiological impairments caused by exposure to tau oligomers, suggesting that manipulating the activity of these enzymes may represent viable therapeutic strategies for combating tau-related impairments. The notion that LCMT-1 or PME-1 dependent increases in PP2A activity may be therapeutically beneficial is supported by published studies of the effects of direct PP2A activators in animal models. Sodium selenate is one such direct PP2A activator that was shown to improve cognitive function and synaptic plasticity in tauopathy models ^31,32,89^, and also reduce tau phosphorylation and improve cognitive function in rodent models of TBI ^33,90,91^. Another promising class of PP2A activators are the SMAPs (small molecule activators of PP2A). SMAPs bind to and stabilize PP2A heterotrimers ^92^. Treatment with one SMAP compound was shown to reduce tau phosphorylation and cognitive impairments in a hyperhomocysteinemic mouse model of AD ^93^. Given the beneficial effects of these compounds in different neurodegenerative animal models, it would be interesting to determine whether they reduce sensitivity to tau-induced impairments in a manner similar to the effects of transgenic LCMT-1 overexpression in this study.

In our experiments, we found that LCMT-1 and PME-1 have opposing effects on tau sensitivity. Given the effect of these enzymes on tau sensitivity, and the reciprocal relationship between them, inhibiting PME-1 might then be expected to lead to decreased tau sensitivity. Eicosanoyl-5-hydroxytryptamide (EHT) is a compound found in coffee that was reported to inhibit PME-1 and increase PP2A activity ^94,95^. EHT administration reduced tau phosphorylation and behavioral impairments caused by overexpression of the PP2A inhibitor, SET/I_2_ PP2A, in rats ^29^. In addition, a small molecule covalent inhibitor of PME-1, aza-beta-lactam 127 (ABL-127), was shown to reduce PP2A demethylation, tau phosphorylation, and oxidative stress in an *in vitro* model of manganese-induced neurotoxicity ^96^. We found previously that treatment with EHT protected mice from behavioral and electrophysiological deficits caused by exposure to soluble oligomers of Aβ ^97^, and EHT treatment was also found to be efficacious in cell and animal models of α-synucleinopathies ^94,98,99^. If EHT, or another PME-1 inhibitor acted similarly to protect against impairments caused by exposure to soluble oligomeric tau, then this would suggest potential therapeutic benefits for PME-1 inhibition in the context of a variety of neurodegenerative conditions with pathogenic mechanisms involving tau, Aβ, and/or α-synuclein.

## METHODS

### Experimental Design

Animals subjected to shock-wave or sham exposure were trained on an accelerating rotarod task for 5 days prior to treatment. Righting reflex was assessed immediately after exposure, and testing on a battery of behavioral tasks was initiated either 13 days or 3 months after exposure. Separate groups of animals were used for testing at each time point, and tissue harvested for western blotting or immunohistochemistry after testing was completed. The behavioral assessments consisted of the following tests conducted in the order listed: open field, accelerating rotarod, elevated plus maze, forced swim test, contextual fear conditioning exposure, radial arm water maze, tail suspension, sensory threshold assessment, and visible platform water maze task. Behavioral experiments conducted on animals infused with tau preparations or vehicle consisted of the tests in the following order: open field testing, radial arm water maze, contextual fear conditioning, visible platform water maze, and sensory threshold assessment.

### Animals

Subjects in this study were 4-6 month-old wild-type mice, or mice carrying combinations of the CaMKII-tTA, tetO-LCMT, and tetO-PME transgenes as indicated. Single transgenic animals carrying either a CaMKII-tTA, tetO-LCMT, or tetO-PME transgene alone were used as controls. Shock-wave exposed animals used in behavioral histological and western blot experiments were generated from crosses of either tetO-LCMT; CaMK-tTA or tetO-PME; CaMK-tTA double transgenic mice in a C57BL6/J background to wild-type mice in a 129S6 background. Animals used for behavioral histological and western blot experiments were generated from crosses of either tetO-LCMT; CaMK-tTA or tetO-PME; CaMK-tTA double transgenic mice in a C57BL6/J background to wild-type C57BL6/J mice. Endogenous murine tau was isolated from shock-wave or sham exposed 3-4 month-old wild-type C57BL6/J mice. Equivalent numbers of male and female mice were used for all experiments and behavioral testing was conducted in groups of 10-12 animals. All procedures involving animals were conducted in strict accordance with protocols approved by the Columbia University Institutional Animal Care and Use Committee (USDA Registration #21-R-0082; AAALAC Accreditation #000687; NYDOH #A141).

### Shockwave exposure

An aluminum shock tube was used in these experiments to deliver high-pressure shockwaves to mice as described previously (Fig. 2A)^61,62^. The shock tube consisted of a 76 mm diameter x 1240 mm long driven section separated from a 25 mm long, helium-pressurized driver section by polyethylene terephthalate membranes. Mice were anesthetized by inhalation of 5% isoflurane/5% oxygen for 2 min followed by 2% isoflurane/5% oxygen for 3 min. Anesthetized mice were placed in a rigid holder that protected the body and internal organs, leaving only the head exposed to the shockwave. The head was positioned at the center of the exit of the shock tube, with the dorsal surface facing the axis of shockwave propagation. The head was supported by a rigid support that extended under its ventral surface, and a nose bar that limited shockwave related movement^101^. Two piezoresistive pressure transducers were flush mounted at the shock tube exit to record shockwave pressure, duration, and impulse. A third pressure transducer was mounted to the interior of the animal holder to detect any pressure changes experienced by the body of the animal. Subcutaneous injections of 5 mg/kg carprofen were administered to mice prior to, 24 hrs, and 48 hrs after shockwave exposure for analgesia. Sham exposed animals were treated identically to shockwave exposed animals with the exception of shockwave exposure. The parameters of the blast injury level were 272 ± 3 kPa peak overpressure, 0.82 ± 0.01 ms duration, and 66 ± 1 kPa*ms impulse (n=12, mean ± SEM).

### Righting time

Immediately following shockwave exposure or sham treatment, anesthetized animals were placed on their backs on an absorbent pad, and the time for all 4 paws to contact the ground was recorded.

### Preparations of tau

#### Recombinant human tau preparation

Recombinant human tau was prepared as described previously^3,64^. Human 4R/2N tau carrying a C-terminal 6x His tag was expressed in *Escherichia coli* from a pET29a vector (gift of Dr. Yoshiaki Furukawa; University of Yokohama, Japan). Pelleted cells were lysed by sonication in buffer containing 2% Triton X-100, 50 mM Na phosphate pH 7.0, 2 mM MgCl_2_ and protease inhibitors (cOmplete Ultra, Roche). Streptomycin sulfate was added to homogenates to precipitate nucleic acids and perchloric acid was added to a final concentration of 0.1%. Homogenates were boiled for 15 min followed by centrifugation at 15,000 x g for 20 min. Supernatants were then neutralized by addition of sodium hydroxide dialyzed against 50 mM NaPO_4_, 300 mM NaCl, 5 mM imidazole, and 0.03% Triton X-100 -pH7.7. His-tagged tau was purified on nickel-agarose spin columns (His-Spin, Zymo Research) and eluted in buffer containing 50 mM NaPO_4_, 300 mM NaCl and 250 mM imidazole – pH 7.7. Eluted tau was then buffer exchanged into 200 mM NaCl, 50 mM Tris-HCl pH 7.2 via Amicon Ultra Centrifugal Devices (Millipore). Final protein concentration was determined by measuring absorption at 280 nm on a Nanodrop spectrophotometer (Thermo Scientific), and oligomerized by incubation in 1 mM hydrogen peroxide for 20 hrs at room temperature.

#### Isolation and enrichment of endogenous murine tau

Endogenous murine tau was isolated as described previously^3,64^. Shockwave or sham exposed mice were euthanized by cervical dislocation. Forebrains were dissected and snap frozen in liquid nitrogen. Samples were stored at -80°C prior to homogenization in ice cold lysis buffer containing 1% perchloric acid, 20 mM L-histidine, and protease (Roche) as well as phosphatase inhibitors (Pierce). For exogenous administration of tau in behavior and electrophysiology experiments, forebrains from ten mice were combined within each group before homogenization. For sELISA analysis, forebrains from four mice were combined within each group before homogenization. After centrifugation at 15,000 *x g*, overnight buffer exchange to 20 mM L-histidine and protein concentration, tau was further enriched by fast protein liquid chromatography (FPLC) using a HiTrap Q HP anion exchange column on an AKTA FPLC device (GE Healthcare). Tau containing fractions were identified by western blot and pooled. The samples were oligomerized via incubation in 1 mM hydrogen peroxide for 20 hrs at room temperature with rotation. For tau de-oligomerization experiments, tau was treated with 5 mM dithiothreitol (DTT) and 5 mM EDTA for 30 min at room temperature with rotation followed by 70°C heat treatment for 10 min.

### Surgical implantation of cannulae and infusion of tau

For experiments involving *in vivo* infusion of tau, animals were implanted with bilateral 26-gauge guide cannulae (Plastics One, Roanoke, VA) into the dorsal part of the hippocampi (coordinates: P=2.46 mm, L=1.50 mm to a depth of 1.30 mm)^102^ under anaesthesia with 20 mg/kg Avertin. Cannulas were fixed to the skull with acrylic dental cement (Paladur) and animals were allowed to recover for 6-8 days following surgery before behavioural testing. One μl of the indicated tau preparation or vehicle was infused at concentrations and timepoints detailed below into each hippocampus over a period of 1 minute using a microsyringe connected to the cannulae by polyethylene tubing. After infusion, the tubing was kept in place for an additional minute to allow diffusion of the injected material into the tissue.

For recombinant tau experiments, LCMT-1 overexpressing and sibling transgenic control mice were infused with 14.85 μg/mL of tau for behavior testing and acute hippocampal slices were perfused with 2.48 μg/mL of tau for electrophysiology. PME-1 overexpressing mice and their transgenic sibling controls were treated with a subthreshold dose of tau that did not produce deleterious effects in control mice. These mice were infused with 4.95 μg/mL of tau for behavior testing and acute hippocampal slices were perfused with 49.5 ng/mL of tau for electrophysiology. For sham and blast tau experiments, wildtype, LCMT-1 overexpressing, and sibling transgenic control mice were infused at a concentration of 22.9 μg/mL for behavior testing and acute hippocampal slices were perfused at a concentration of 114.7 ng/mL for electrophysiology. PME-1 overexpressing mice and their transgenic sibling controls were treated at a concentration of 4.59 μg/mL for behavior testing and acute hippocampal slices were perfused at a concentration of 22.9 ng/mL.

### Behavior

#### Open field behavior

Sham or shockwave exposed animals were subjected to a novel open field environment as previously described^49^. Mice were placed in a plexiglass chamber (27.3 cm long × 27.3 cm wide × 20.3 cm high) (model ENV-510, Med Associates) for a total of 20 min during which time their movements were tracked and analyzed using an overhead video tracking system and behavioral analysis software (Ethovision XT, Noldus). Animals infused with tau or vehicle were placed in a similar plexiglass arena housed in a sound attenuating chamber for 10 min on each of two successive days during which time their movements were tracked using arrays of infrared beams connected to a computerized tracking system, and analyzed using behavioral analysis software (Activity Monitor, Med Associates). For infusion experiments tau preparations or vehicle was administered at 180 and 20 min prior to testing on each day. In all experiments, the “center” of the open field was defined as an area beginning 10 cm from the walls of the arena.

#### 2-day radial arm water maze (RAWM) task

Testing was performed in a 120 cm diameter pool containing a six-arm radial maze insert (San Diego Instruments) and filled with opaque water as described previously^50^. Mice were tested in 15 x 1-minute trials on each of 2 consecutive days. The location of the escape platform was held constant during testing but the start location was pseudorandomly varied throughout. On the first day, training alternated between visible and hidden platform trials, while on the second day only hidden platform trials were conducted. Water temperature was maintained at approximately 24° C and mice were dried and placed in a clean heated cage between trials to prevent hypothermia. Entries into maze arms that did not contain the escape platform were scored as errors. Data are presented as the average number of errors committed during blocks of 3 training trials. Animals that received tau preparations or vehicle treatment were infused 180 and 20 min prior to the start of testing on each day and again 180 and 20 min prior to the start of the seventh trial of each day.

#### Visible platform water maze task

Two versions of the visible platform water maze task were performed. In each case the apparatus consisted of a 120 cm diameter pool (San Diego Instruments) filled with opaque water maintained at approximately 24° C as described previously^49^. Trial duration was a maximum of 120 sec during which time animals were required to swim to a marked escape platform located at the water surface. Intertrial intervals during trial blocks were 15 -20 min, during which time animals were dried briefly with paper towels before being placed in their home cage. Animals that did not reach the platform within the allotted time were guided to it and allowed to sit there for 15 sec. Platform locations were changed between trials such that it was not present in the same quadrant on any two successive trials. Start locations also alternated among the quadrants adjacent and opposite to the visible platform. Animals that were subjected to shockwave or sham exposure were tested during 4 trials on each of two successive days, and movements were recorded and analyzed using a video-tracking system and behavioral analysis software (ANY-maze). Animals infused with tau preparations or vehicle were tested during 3 morning and 3 afternoon trials on each of two successive days, and movements were recorded and analyzed using a video-tracking system behavioral analysis software (Ethovision XT, Noldus). Intrahippocampal infusions of tau or vehicle were performed 180 and 20 min prior to the start of each block of 3 trials.

#### Contextual fear conditioning

As previously described^49^, animals were placed into a transparent Plexiglas conditioning chamber (33cm x 20cm x 22cm) (Noldus PhenoTyper). Animal movement was recorded using an overhead video camera connected to a personal computer and freezing behavior was scored and analyzed using Ethovision XT software (Noldus). Foot shocks were administered through a removable metal grid floor and the entire apparatus was cleaned and deodorized between animals with distilled water and 70% ethanol. Animals were placed in the conditioning chamber once on each of two consecutive days. On the first day of exposure mice were placed in the conditioning chamber for 2 minutes before the onset of a discrete 30s, 2800Hz, 85dB tone, the last 2s of which coincided with a 0.8 mA foot shock. After the tone and shock exposure, the mice were left in the conditioning chamber for another 30s before returning to their home cages. 24 hrs after their first exposure, animals were returned to the conditioning chamber for 5 min without foot shock or tone presentation. Animals that received tau preparations or vehicle treatment were infused 180 and 20 min prior to exposure on the first day only.

#### Sensory threshold assessment

As previously described^49^, animals were placed into the same apparatus used for contextual fear conditioning. A sequence of single, 1 sec foot shocks were then administered at 30 sec intervals and 0.1 mA increments from 0 up to a maximum of 0.7 mA. Each animal’s behavior was monitored by the experimenter to determine their thresholds for first visible response to the shock (flinch), their first gross motor response (run/jump), and their first vocalized response. Animals that received tau preparations or vehicle treatment were infused 180 and 20 min prior to testing.

#### Accelerating rotarod task

We assessed motor performance of mice using a rotarod apparatus (Rota-Rod, Letica Scientific Instruments) essentially as described previously^103,104^ (Wang, 2011 #126). Training on this task was carried out on 5 successive days. The first day of consisted of pretraining carried out over 3 x 3 min trials. During the first two of these trials, animals were placed on the apparatus and allowed to habituate for 3 minutes, animals that fell from the stationary rod were replaced until the trial was completed. During the third trial, the rotation speed was fixed at 4 rpm and animals that fell from the rod were also replaced until the trial was completed. Training on all subsequent days consisted of 3 × 5 min trials per day with the rotation speed ramped from 4 to 22 rpm over the course of the trial. When animals fell from the apparatus the trial was terminated and the animal returned to its home cage. Testing at 2 weeks or 3 months after shockwave exposure consisted of 3 trials conducted in the same manner as described for training days 2-5. The intertrial interval at all stages of the experiment was 15 - 20 min.

#### Elevated plus maze

Testing in an elevated plus maze was performed essentially as previously described^105,106^. The apparatus consists of a plus shaped track with arms 35 cm long and 5 cm wide, elevated 50 cm above the bench top (Model 60410, Stoelting). Two non-adjacent arms are surrounded by 15 cm high walls on 3 sides, and the remaining two arms were open. Animals were placed into the center of the apparatus and their location during single 6 min exposure to this apparatus was monitored and analyzed using a video tracking system and accompanying behavioral analysis software (ANY-maze).

#### Forced swim test

Animals were tested in a forced swim test essentially as previously described^107,108^. Animals were placed into a 4-liter glass beaker filled half way with tap water (22-25 °C) for a total of 6 min. During this time, the animals’ movements were recorded using a video camera, and periods of immobility were scored offline by an observer blinded to genotype and treatment.

#### Tail suspension test

Animals were tested in a tail suspension test essentially as previously described^105^. Animals’ tails were gently taped approximately 2 cm from the end to a horizontal bar elevated 30 cm above the benchtop. The animals were then suspended in this position for 6 minutes while their movements were recorded using a digital video camera. Periods of immobility were then scored offline by an observer blinded to genotype and treatment.

### Electrophysiological studies

Extracellular field potential recordings were performed on acute hippocampal slices prepared as described previously^3^. Animals were euthanized by cervical dislocation, and brains were removed rapidly and cooled in ice cold ASCF consisting of in mM: 124 NaCl, 4.4 KCl, 1 Na_2_HPO_4_, 25 NaHCO_3_, 2 CaCl_2_, 2 MgCl_2_, and 10 glucose. Hippocampi were then dissected and sliced into 400 μM sections using a tissue chopper. Slices were incubated at 29°C in an interface chamber under continuous perfusion (2 ml/min) with oxygenated ACSF and allowed to recover for a minimum of 90 min prior to recording responses in the CA1 region to stimulation of Schaffer collateral projections with a bipolar electrode (FHC Inc.). Field potential signals were acquired using an Axoclamp-2A amplifier (Axon Instruments) and pClamp10.6 software (Molecular Devices). Input/output relationships were determined prior to each recording and stimulus intensities that elicited 30% of the maximal response were utilized. Stable baselines were obtained for a minimum of 15 min prior to application of tau or vehicle. LTP was elicited by application of a theta-burst stimulation protocol consisting of 3 trains separated by 15 second intervals with each train consisting of 10 bursts at 5 Hz and each burst consisting of 5 pulses at 100 Hz. Data analysis was performed using Clampfit 10.6 software (Molecular Devices).

### Western blotting

Animals were euthanized by cervical dislocation. Hippocampi were then rapidly dissected, snap frozen in liquid nitrogen and stored at -80°C prior to homogenization for western blot analysis. Hippocampal homogenates were prepared by sonication at 95°C in aqueous buffer containing 2% lithium dodecyl sulfate and 50 mM Tris pH 7.5. Total protein concentrations were determined by bicinchoninic acid assay according to the manufacturer’s instructions (Micro BCA protein assay kit, Thermo Fisher) and 20 μg of total protein was loaded per lane on NuPage 4-12% Bis-Tris gels (Invitrogen). Proteins were transferred to PVDF membranes using an i-Blot gel transfer device (Invitrogen). Membranes were blocked with Seablock (Pierce) for 1 hr at room temperature, and probed overnight at 4°C with the indicated primary antibodies: mouse anti-phospho-tau clone PHF1 and clone CP13 diluted 1:500 (gifts from P. Davies, Feinstein Institutes); mouse anti-phospho-tau clone AT270 diluted 1:1000 (Thermo fisher #MN1050 RRID:AB_223651), rabbit anti-tau clone EP2456Y diluted 1:8000 (Abcam #Ab76128 RRID:AB_1524475) mouse anti-β-actin diluted 1:40,000 (Licor #926-42212, RRID:AB_2756372); rabbit anti-β-actin diluted 1:40,000 (Licor #926-42210, RRID:AB_1850027). Membranes were then washed and incubated with infrared dye-labeled Goat anti-rabbit (IRDye 800CW, LI-COR) and Goat anti-mouse (IRDye 680RD, LI-COR) secondary antibodies at room temperature for 2 hrs. Immunoreactive bands were detected using an Odyssey 9120 infrared imaging system and analyzed using Image Studio Lite v5.2.5 software (RRID:SCR_014211) (LI-COR).

### Sandwich enzyme linked immunosorbent assay (sELISA)

sELISA for total tau was performed using the mouse total tau sELISA kit (ThermoFisher catalog KMB7011) according to the manufacturer’s instructions. sELISA for oligomeric tau was performed as previously described^3^. TOC1 antibody (tau oligomers) was used for capture and R1 antibody (polyclonal rabbit tau antibody) was used for detection of bound tau. All steps in this assay were performed at room temperature. TOC1 was diluted to 2 μg/ml in borate saline (100 mM boric acid, 25 mM sodium tetraborate decahydrate, 75 mM NaCl, 250 μM thimerosal) and incubated in a high binding sELISA microplate (Corning, #3590) for 1 hour. The sELISA plate was washed twice with 200 μl/well of a buffer containing 100 mM boric acid, 25 mM sodium tetraborate decahydrate, 75 mM NaCl, 250 μM thimerosal, 0.4% bovine serum albumin and 0.1% tween-20. The plate was then incubated with 200 μl/well of a blocking buffer consisting of 5% non-fat milk for 1 hour. The plate was rinsed twice with the wash buffer and then samples were added to the wells for 1.5 hours. R1 antibody was diluted to 0.1 μg/ml in the blocking buffer. Wells were rinsed twice, incubated with R1 for 1.5 hours and rinsed another 3 times. Wells were then incubated with goat anti-rabbit antibody conjugated to horseradish peroxidase (Vector Labs, PI-1000), which was diluted to 0.2 μg/mL in blocking reagent, for 1.5 hours and then rinsed 3 times. The signal was developed by incubation with 3,3⍰,5,5⍰-tetramethylbenzidine (TMB) for 10-15 min and the reaction was stopped using 3.5% sulfuric acid. The TMB signal was measured at 450 nm absorbance wavelength.

### Statistical analysis

Experiments were performed blind, and results are shown as mean ± SEM. Tests for statistical significance between groups were performed using Prism 9 (Graphpad Software, San Diego, CA, USA). Student’s unpaired, one or two-tailed t-tests as indicated were used for experiments involving 2 groups. 2-way ANOVA comparisons – with or without repeated measures as dictated by the experimental design – were used for analysis of all experiments involving multiple groups. Post-hoc comparisons for multiple group experiments were performed using Dunnett’s tests when comparisons were made to a single control group, and Sidak’s post-hoc tests were performed on data from experiments without repeated measures. Tukey’s tests were used when comparisons were made among multiple groups.

## Acknowledgements

We would like to thank the late Dr. Peter Davies for the gift of the PHF-1 and CP13 antibodies. We acknowledge Tessa Grabinski for her execution of the TOC1 sELISA. We also would like to acknowledge Dr. Anthony Pacifico (Department of Veteran Affairs; the views expressed are those of the authors and do not necessarily represent the views of the U.S. Department of Veteran Affairs or other U.S. Government entity) for valuable discussions. This work was supported by the Department of Defense grant (W81XWH-12-1-0579, W81XWH-15-1-0550, W81XWH-21-1-0359, HT9425-23-1-0384) and the Paul G. Allen Frontiers group, grant 12347.

## Author contribution

Conceptualization: REN, OA, BM

Experimentation: SS, REN, EWV, CDH, HZ, AS, EA, ZG, KA, SL, LS, MG, HLB, MF, NMK

Data analysis: SS, REN, OA, NK, EWV, NMK

Supervision: REN, OA, BM

Writing: SS, REN

Editing manuscript: OA, REN, BM, SS, NMK

