## Supplementary Figures for "PP2A methylesterase, PME-1, and PP2A methyltransferase, LCMT-1, control sensitivity to impairments caused by injury-related oligomeric tau"

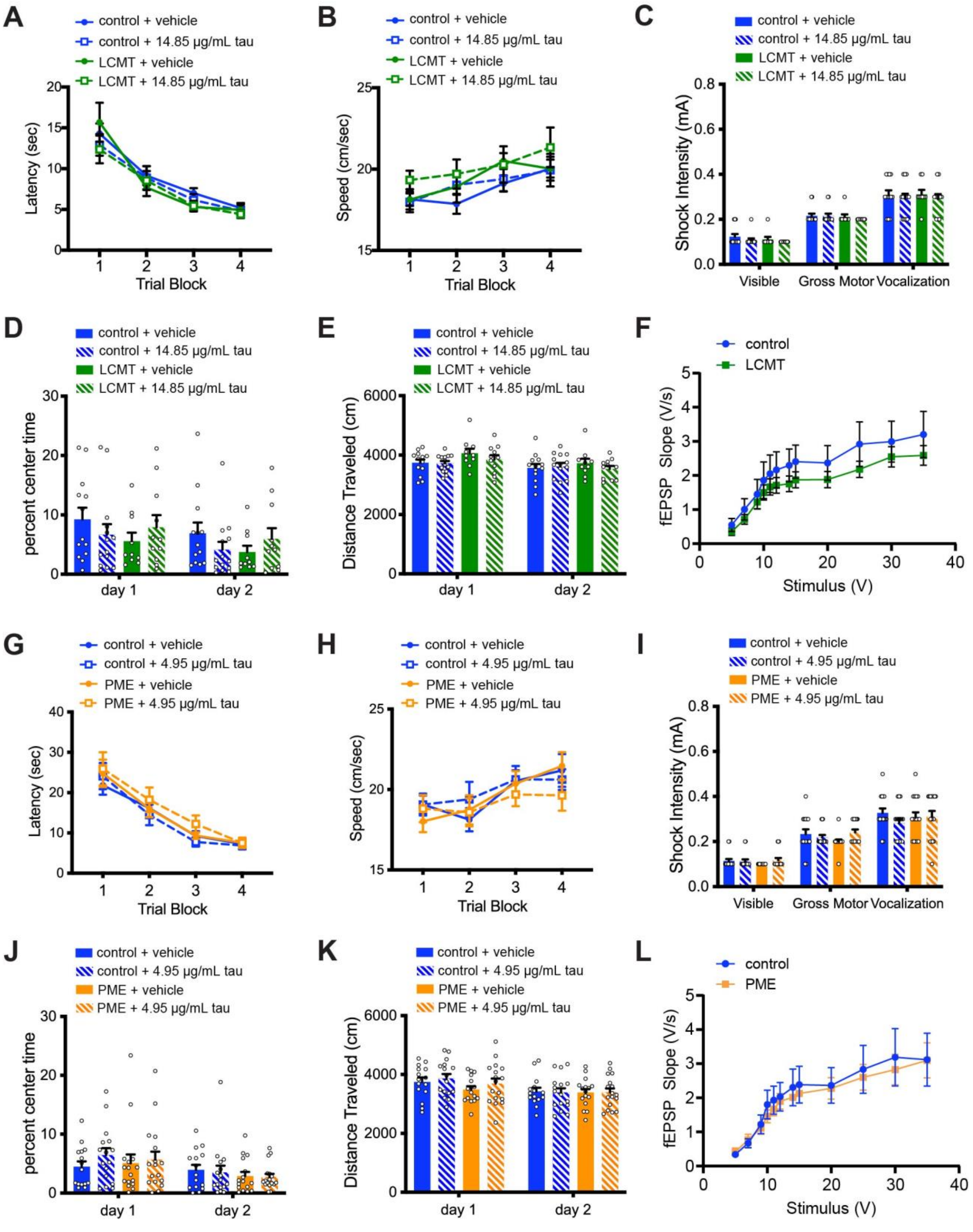

**Fig. S1: Transgenic overexpression of LCMT-1 or PME-1 does not alter the open field behavior, foot shock perception, or marked platform water maze performance of mice treated with recombinant oligomeric tau.** **A)** Time to reach the escape platform in a marked platform version of the Morris water maze task conducted in 4 blocks of 3 trials over two days for LCMT-1 overexpressing transgenic mice and sibling controls infused with recombinant oligomeric tau or vehicle shows no significant differences among groups (2-way RM ANOVA for latency with group and trial as factors shows no significant effect of group ($F(3,47) = 0.5326$, $P = 0.6622$), but a significant effect of trial ($F(3,141) = 62.01$, $P < 0.0001$). **B)** Swim speed for the indicated groups during the marked platform version of the Morris water maze task shown in (A). 2-way RM ANOVA for speed with group and trial as factors shows no significant effect of group ($F(3,47) = 1.133$, $P = 0.3453$), but a significant effect of trial ($F(3,141) = 8.091$, $P < 0.0001$). **C)** Threshold for responses to foot shocks of increasing intensity for LCMT-1 overexpressing transgenic mice and sibling controls infused with recombinant oligomeric tau or vehicle shows no significant differences among groups (2-way RM ANOVA for effect of group $F(3,42) = 0.6673$, $P = 0.5769$; Tukey's post-hoc comparisons revealed no significant differences between treatment groups for any response, $P > 0.05$). **D)** Time spent in the center of an open field environment during two 10-minute trials over two days for LCMT-1 overexpressing transgenic mice and sibling controls infused with recombinant oligomeric tau or vehicle shows no significant differences among treatment groups (2-way RM ANOVA for effect of group: $F(3,44) = 1.257$, $P = 0.3008$). Moreover, Tukey's multiple comparisons show no significant differences between any groups on either day ($P > 0.05$). N = 13 control+vehicle, 14 control+tau, 10 LCMT+vehicle, 11 LCMT+tau mice. **E)** Graph shows no significant difference in distance traveled by the same animals in (D) during two 10 minute trials over two days in an open field environment (2-way ANOVA for effect of group: $F(3,44) = 0.9956$, $P = 0.4038$; Tukey's post-hoc pairwise comparisons show no significant differences between groups on either day ($P > 0.05$.). **F)** Stimulus-response relationship at Schaffer collateral synapses in hippocampal slices prepared from LCMT overexpressing or transgenic control mice before treatment with tau shows no significant differences between groups (2-way RM ANOVA $F(1,56) = 0.6971$, $P = 0.4073$). **G)** Time to reach the escape platform in a marked platform version of the Morris water maze task for PME-1 overexpressing transgenic mice and sibling controls infused with recombinant oligomeric tau or vehicle shows no significant differences among groups (2-way RM ANOVA for latency with group and trial as factors shows no significant effect of group ($F(3,60) = 0.5564$, $P = 0.6459$), but a significant effect of trial ($F(3,180) = 66.69$, $P < 0.0001$). **H)** Swim speed for the indicated groups during the marked platform version of the Morris water maze task shown in (G). 2-way RM ANOVA for speed with group and trial as factors shows no significant effect of group ($F(3,60) = 0.2641$, $P = 0.8510$), but a significant effect of trial ($F(3,180) = 4.300$, $P = 0.0001$). **I)** Threshold for responses to foot shocks of increasing intensity for PME-1 overexpressing transgenic mice and sibling controls infused with recombinant oligomeric tau or vehicle shows no significant differences among groups (2-way RM ANOVA for effect of group $F(3,60) = 1.322$, $P = 0.2758$; Tukey's post-hoc comparisons revealed no significant differences between treatment groups for any response, $P > 0.05$). **J)** Time spent in the center of an open field environment during two 10 minute trials over two days for PME-1 overexpressing transgenic mice and sibling controls infused with recombinant oligomeric tau or vehicle shows no significant differences among treatment groups (2-way RM ANOVA for effect of group: $F(3,60) = 0.2471$, $P = 0.8631$). Moreover, Tukey's multiple comparisons show no significant differences between any groups on either day ($P > 0.05$). N = 15 vehicle, 16 control+tau, 16 PME+vehicle, 17 PME+tau mice. **K)** Graph shows no significant difference in distance traveled by the same animals in (J) during two 10 minute trials over two days in an open field environment (2-way ANOVA for effect of group: $F(3,60) = 0.4761$, $P = 0.7001$; Tukey's post-hoc pairwise comparisons show no significant differences between groups on either day ($P > 0.05$.). **L)** Stimulus-response relationship at Schaffer collateral synapses in hippocampal slices prepared from PME overexpressing or transgenic control mice before treatment with tau shows no significant differences between groups (2-way RM ANOVA $F(1,38) = 0.08794$, $P = 0.7684$). All data presented as mean $\pm$ SEM.

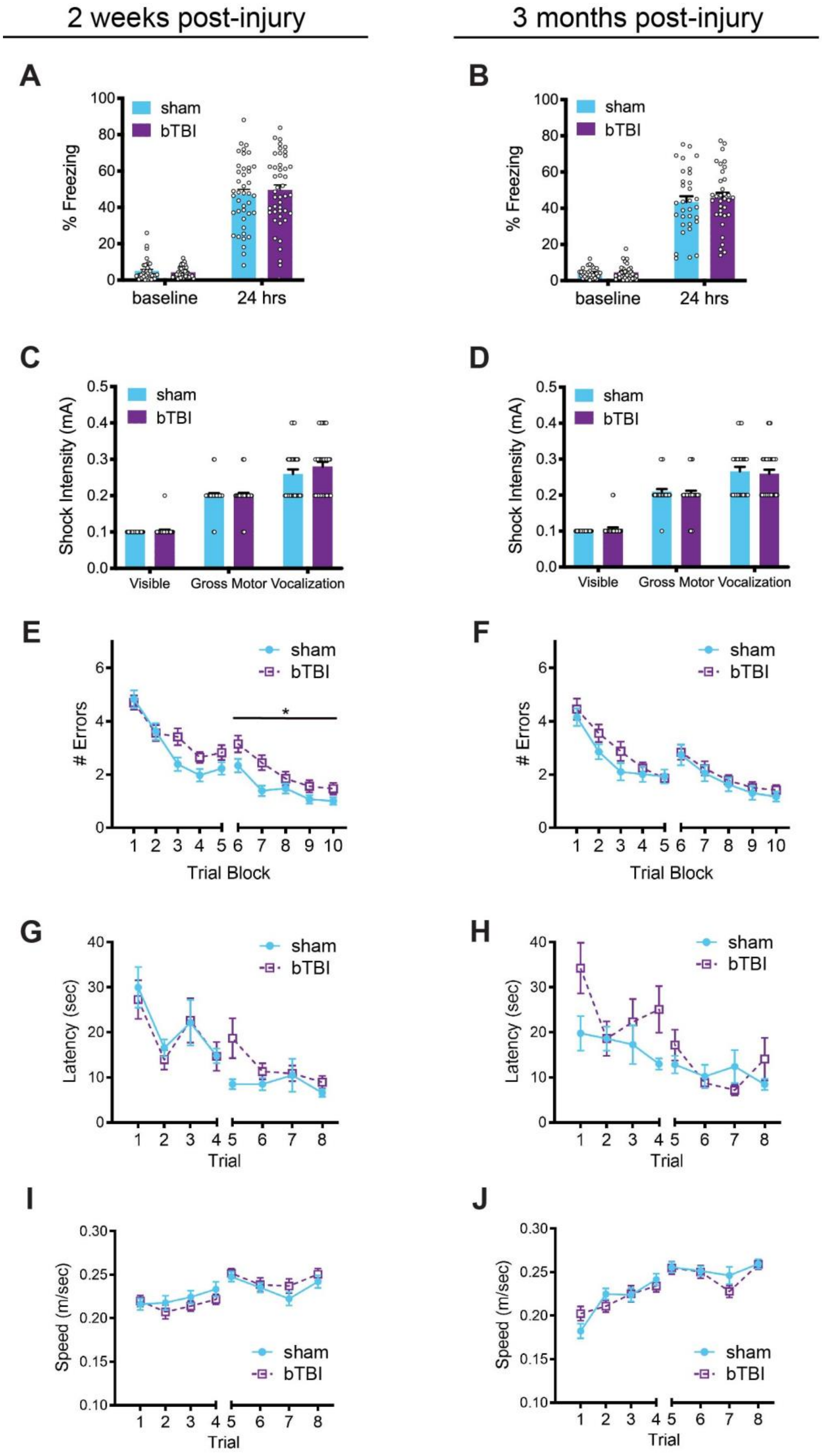
2 weeks post-injury
3 months post-injury
A
B
% Freezing
sham
bTBI
baseline
24 hrs
C
D
Shock Intensity (mA)
Visible
Gross Motor
Vocalization
E
F
### Errors
Trial Block
*
G
H
Latency (sec)
Trial
I
J
Speed (m/sec)
Trial

**Fig. S2: Shockwave exposure does not significantly affect learning and memory performance in a contextual fear condition task, but does result in small but significant impairment in a 2-day RAWM task at 2 weeks but not 3 months after injury. A)** Graph shows no significant differences in time spent freezing during an initial 3 min pre-shock exposure (baseline) to a contextual fear conditioning apparatus (unpaired, two-tailed t-test: t = 0.9331, P = 0.3535), or during subsequent 5 min post-shock exposure the next day (24 hrs) (unpaired, two-tailed t-test t = 0.6178, P = 0.5384) for sham and shockwave (bTBI) exposed animals 15 days after injury (N = 41 sham, 44 bTBI). **B)** Graph shows no significant differences in time spent freezing during an initial 3 min pre-shock exposure (baseline) to a contextual fear conditioning apparatus (unpaired, two-tailed t-test: t = 0.2544, P = 0.8000), or during subsequent 5 min post-shock exposure the next day (24 hrs) (unpaired, two-tailed t-test t = 0.5395, P = 0.5914) for sham and shockwave exposed animals 3 months after injury (N = 32 sham, 34 bTBI). **C)** Threshold for responses to foot shocks of increasing intensity for sham and shockwave (bTBI) exposed animals 20 days after injury revealed no significant differences in foot-shock perception (2-way RM ANOVA for effect of treatment F(1,64) = 0.8890, P = 0.3493; for response F(2,128) = 237, P < 0.0001, Sidak post-hoc comparisons revealed no significant differences between treatment groups for any response, P > 0.05; N=32 sham, 34 bTBI for this and the following panels). **D)** Threshold for responses to foot shocks of increasing intensity for sham and shockwave exposed animals 3 months after injury revealed no significant differences in foot-shock perception (2-way RM ANOVA for effect of treatment F(1,64) = 0.8890, P = 0.3493; for response F(2,128) = 237, P < 0.0001, Sidak post-hoc comparisons revealed no significant differences between treatment groups for any response, P > 0.05). **E)** Graph shows a significant difference in number of errors committed during the second day of testing (blocks 6-10) on a 2-day RAWM task for sham and shockwave exposed animals 17 days after injury (2-way RM ANOVA for effect of treatment F(1,87) = 5.405, P = 0.0224; block F(4,348) = 35.17, P < 0.0001; interaction F(4,348) = 1.975, P = 0.0979; N = 43 sham, 46 bTBI). **F)** Graph shows no significant difference in number of errors committed during the second day of testing (blocks 6-10) on a 2-day RAWM task for sham and shockwave exposed animals 3 months after injury (2-way RM ANOVA for effect of treatment F(1,64) = 0.3364, P = 0.5640; block F(4,256) = 27.59, P < 0.0001; interaction F(4,256) = 0.06159, P = 09930; N = 32 sham, 34 bTBI). **G)** Time to reach the escape platform for sham and shockwave exposed animals 22 days after injury during each of 8 trials conducted over two successive days in a marked platform version of the Morris water maze task. 2-way RM ANOVA for latency with treatment and trial as factors shows no significant effect of treatment (F(1,64) = 0.5009, P = 0.481), but a significant effect of trial (F(7,448) = 10.63, P < 0.0001). **H)** Time to reach the escape platform for sham and shockwave exposed animals 3 months after injury during each of 8 trials conducted over two successive days in a marked platform version of the Morris water maze task. 2-way RM ANOVA for latency with treatment and trial as factors shows no significant effect of treatment (F(1,64) = 3.763, P = 0.0568), but a significant effect of trial (F(7,448) = 10.63, P < 0.0001). **I)** Swim speed for sham and shockwave exposed animals 22 days after injury during each of 8 trials conducted over two successive days in a marked platform version of the Morris water maze task shown in (G). 2-way RM ANOVA for speed with treatment and trial as factors shows no significant effect of treatment (F(1,64) = 0.0011, P = 0.9733), but a significant effect of trial (F(7,448) = 8.941, P < 0.0001). **J)** Swim speed for sham and shockwave exposed animals 3 months after injury during each of 8 trials conducted over two successive days in a marked platform version of the Morris water maze task shown in (H). 2-way RM ANOVA for latency with treatment and trial as factors shows no significant effect of treatment (F(1,64) = 0.2033, P = 0.6536), but a significant effect of trial (F(7,448) = 23.95, P < 0.0001). All data presented as mean $\pm$ SEM.

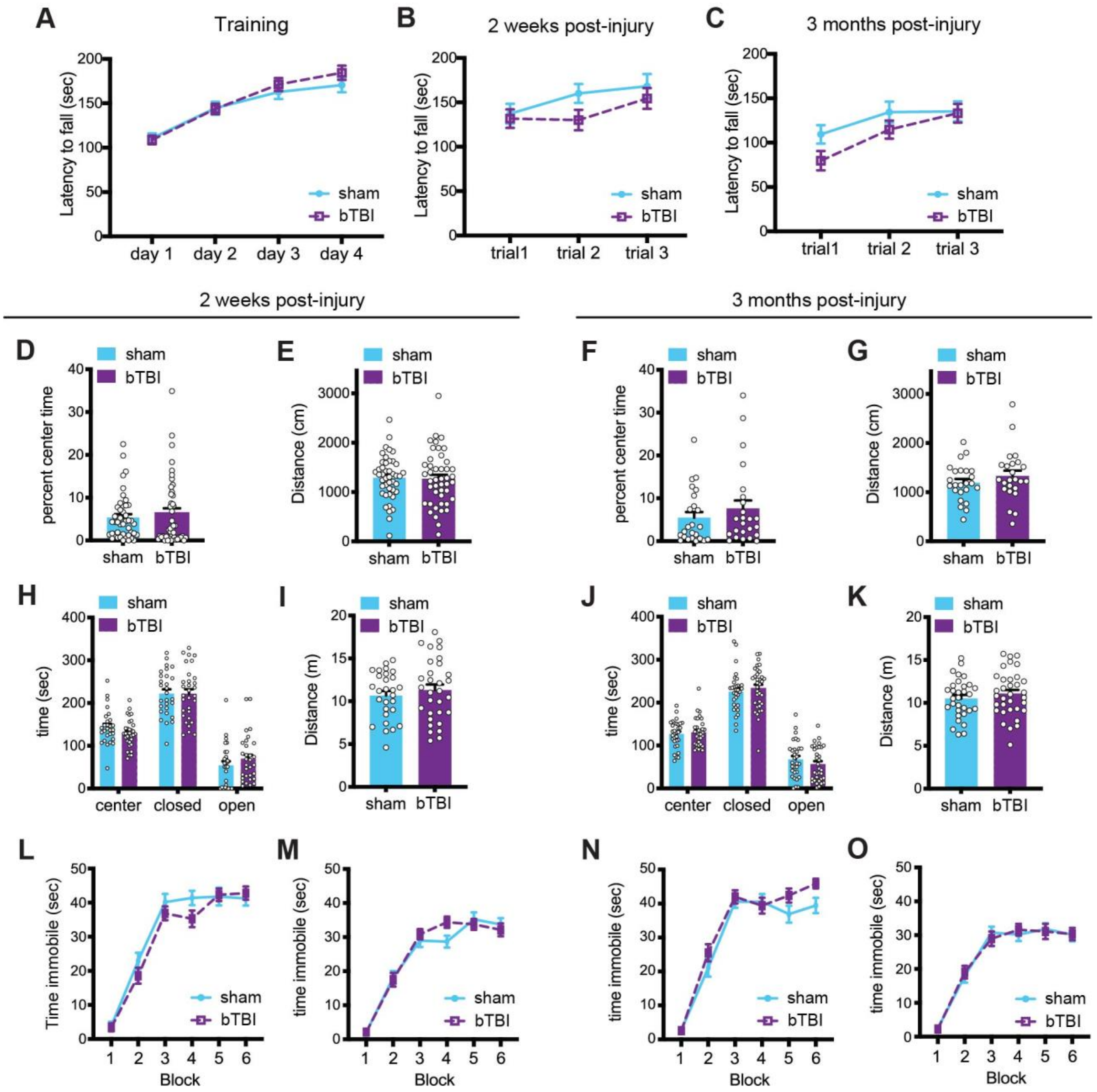


**Fig. S3: Shockwave exposure does not significantly impact motor performance, anxiety, or depression. A)** Latency to fall on an accelerating rotarod task during two training trials on each of 4 days prior to shockwave (bTBI) exposure or sham treatment shows no significant difference between groups but a significant effect of training (2-way RM ANOVA for effect of treatment $F(1,155) = 0.3378$, $P = 0.5619$; day $F(3,465) = 104.5$, $P < 0.0001$, Sidak post-hoc comparisons revealed no significant differences between groups on any day, $P > 0.05$; N = 76 sham, 81 bTBI). **B)** Latency to fall on an accelerating rotarod task during each of three training trials performed 13 days after shockwave or sham exposure shows no significant difference between groups (2-way ANOVA for effect of treatment $F(1,87) = 1.247$, $P = 0.2673$; Sidak post-hoc comparisons revealed no significant differences between groups in any trial, $P > 0.05$; N = 43 sham, 46 bTBI). **C)** Latency to fall on an accelerating rotarod task during each of three training trials performed 3 months after shockwave or sham exposure shows no significant difference between groups (2-way ANOVA for effect of treatment $F(1,64) = 1.474$, $P = 0.2292$; Sidak post-hoc comparisons revealed no significant differences between groups in any trial, $P > 0.05$; N = 32 sham, 34 bTBI). **D)** Time spent in the center of an open field environment during a 20 min trial revealed no significant differences between sham and shockwave exposed animals 13 days after injury (unpaired, one-tailed t test: $t = 0.7201$, $P = 0.2368$; N = 44 sham, 45 bTBI). **E)** Graph shows no significant difference in distance traveled by the same animals in (D) during a 20 min exposure to an open field environment (unpaired, two-tailed t-test: $t = 0.2005$, $P = 0.8415$). **F)** Graph shows no significant difference in time spent in the center of an open field environment during a 20 min trial for sham and shockwave exposed animals 3 months after injury (unpaired, one-tailed t test: $t = 0.9228$, $P = 0.1808$; N = 24 sham, 24 bTBI). **G)** Graph shows no significant difference in distance traveled by the same animals in (F) during a 20 min exposure to an open field environment (unpaired, two-tailed t test: $t = 1.048$, $P = 0.2999$). **H)** Graph shows no significant differences in time spent in the center, closed, or open arms during 6 minutes of exposure to an elevated plus maze for sham and shockwave exposed animals 14 days after injury (2-way ANOVA for effect of treatment $F(1,57) < 0.0001$, $P > 0.9999$; Sidak multiple comparisons showed no significant differences between groups in any zone: center $P = 0.6072$, closed $P > 0.9999$, open $P = 0.5721$; N = 28 sham, 31 bTBI). **I)** Graph shows no significant difference in distance traveled by the same animals in (H) during elevated plus maze testing (unpaired, two-tailed t-test $t = 0.7474$, $P = 0.4579$). **J)** Graph shows no significant differences in time spent in the center, closed, or open arms during 6 minutes of exposure to an elevated plus maze for sham and shockwave exposed animals 3 months after injury (2-way ANOVA for effect of treatment $F(1,63) = 0.29$, $P = 0.5921$; Sidak multiple comparisons showed no significant differences between groups in each zone: center $P = 0.9851$, closed $P = 0.8124$, open $P = 0.6097$; N = 31 sham, 34 bTBI). **K)** Graph shows no significant difference in distance traveled by the same animals in (J) during elevated plus maze testing (unpaired, two-tailed t-test $t = 0.8838$, $P = 0.3801$). **L)** Graph shows no significant difference in time spent immobile during each 1 min block of a 6 min forced swim task for sham and shockwave exposed animals 14 days after injury (2-way RM ANOVA for effect of treatment $F(1,63) = 1.212$, $P = 0.2751$; block $F(5,315) = 149.9$, $P < 0.0001$; Sidak post-hoc comparisons revealed no significant differences between treatment groups during any block, $P > 0.05$; N = 32 sham, 33 bTBI). **M)** Graph shows no significant difference in time spent immobile during each 1 min block of a 6 min tail suspension task for sham or shockwave exposed animals 17 days after injury (2-way RM ANOVA for effect of treatment $F(1,63) = 0.1702$, $P = 0.6813$; block $F(5,315) = 168.1$, $P < 0.0001$; Sidak post-hoc comparisons revealed no significant differences between treatment groups during any block, $P > 0.05$; N = 31 sham, 34 bTBI). **N)** Graph shows no significant difference in time spent immobile during each 1 min block of a 6 min forced swim task for sham and shockwave exposed animals 3 months after injury (2-way RM ANOVA for effect of treatment $F(1,64) = 2.024$, $P = 0.1597$; block $F(5,320) = 179.4$, $P < 0.0001$; Sidak post-hoc comparisons revealed no significant differences between treatment groups during any block, $P > 0.05$; N = 32 sham, 34 bTBI). **O)** Graph shows no significant difference in time spent immobile during each 1 min block of a 6 min tail suspension task for sham and shockwave exposed animals 3 months after injury (2-way RM ANOVA for effect of treatment $F(1,58) < 0.0001$, $P = 0.9995$; block $F(5,290) = 99.18$, $P < 0.0001$; Sidak post-hoc comparisons revealed no significant differences between treatment groups during any block, $P > 0.05$; N = 29 sham, 31 bTBI). All data presented as mean $\pm$ SEM.

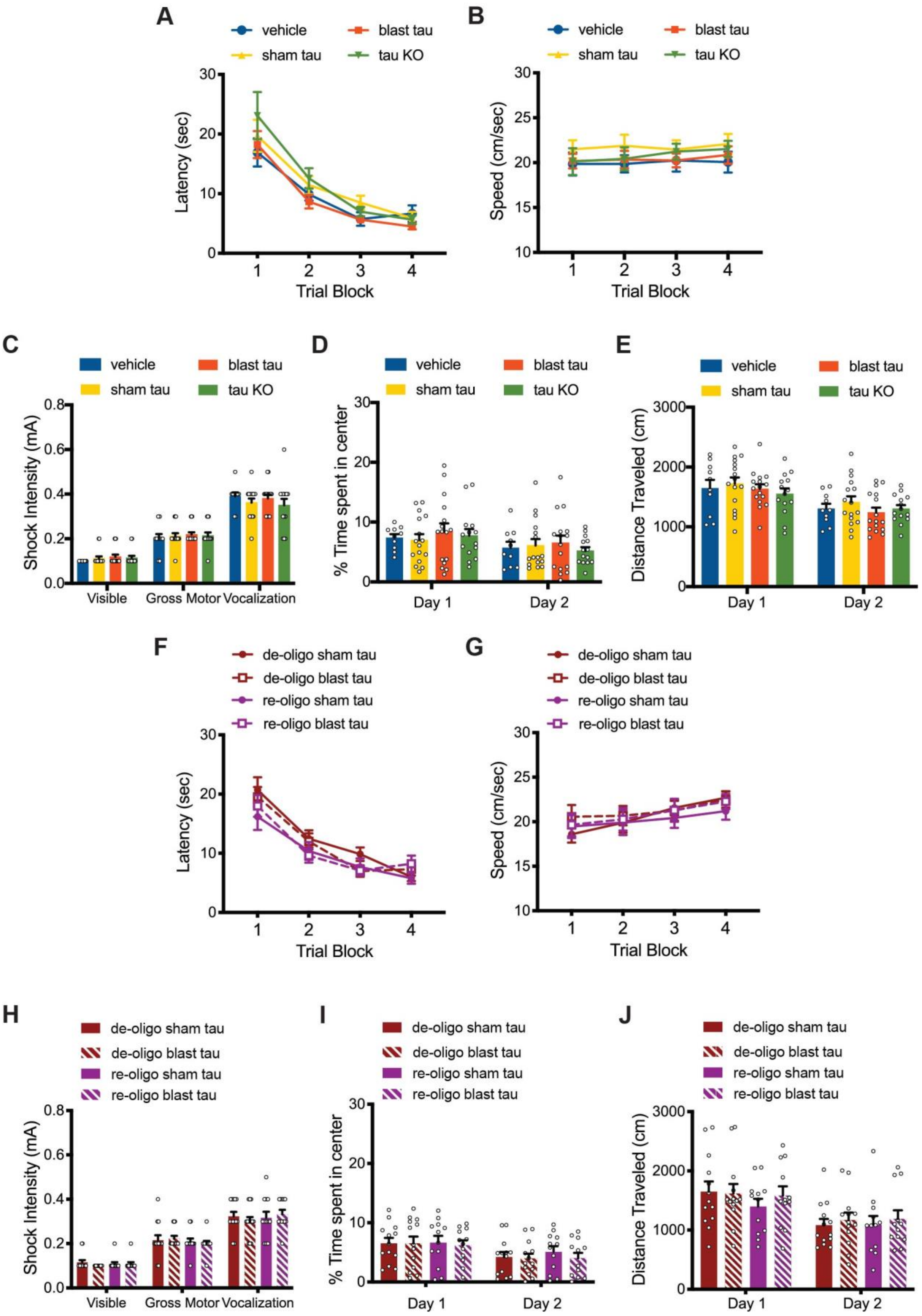
A
vehicle
blast tau
sham tau
tau KO
Latency (sec)
Trial Block
B
Speed (cm/sec)
Trial Block
C
Shock Intensity (mA)
Visible
Gross Motor
Vocalization
D
% Time spent in center
Day 1
Day 2
E
Distance Traveled (cm)
F
de-oligo sham tau
de-oligo blast tau
re-oligo sham tau
re-oligo blast tau
G
H
I
J

**Fig. S4: Oligomerized tau isolated from shockwave exposed mice does not affect open field behavior, foot shock perception, or performance in a marked platform version of the Morris water maze.** **A)** Time to reach the escape platform in a marked platform version of the Morris water maze task conducted in 4 blocks of 3 trials over two days for mice infused with vehicle, sham tau, blast tau, or mock purified material from shockwave exposed tau KO mice shows no significant differences among treatment groups (2-way RM ANOVA for latency with treatment and trial as factors shows no significant effect of treatment ($F(3,52) = 1.238$, $P = 0.3052$), but a significant effect of trial ($F(3,156) = 68.98$, $P < 0.0001$). **B)** Swim speed for the indicated groups during the marked platform version of the Morris water maze task shown in A. 2-way RM ANOVA for speed with treatment and trial as factors shows no significant effects of treatment ($F(3,52) = 0.5991$, $P = 0.6184$) or trial ($F(3,156) = 0.9860$, $P = 0.4010$). **C)** Threshold for responses to foot shocks of increasing intensity for mice infused with vehicle, sham tau, blast tau, or mock purified material from shockwave exposed tau KO mice shows no significant differences among treatment groups (2-way RM ANOVA for effect of treatment $F(3,52) = 0.3384$, $P = 0.7977$; Tukey's post-hoc comparisons revealed no significant differences between treatment groups for any response, $P > 0.05$). **D)** Time spent in the center of an open field environment during a 20-minute trial for mice infused with vehicle, sham tau, blast tau, or mock purified material from shockwave exposed tau KO mice shows no significant differences among groups (2-way RM ANOVA for effect of group: $F(3,44) = 0.2005$, $P = 0.8955$). Moreover, Tukey's multiple comparisons show no significant differences between any groups on either day ($P > 0.05$). N = 10 vehicle, 16 sham tau, 16 sham tau, 14 tau KO. **E)** Graph shows no significant difference in distance traveled by the same animals in (D) during a 20-minute exposure to an open field environment (2-way ANOVA for treatment with treatment and day as factors $F(3,52) = 0.6735$, $P = 0.5722$; Tukey's post-hoc pairwise comparisons show no significant differences between groups on either day ($P > 0.05$.)**.** **F)** Time to reach the escape platform in a marked platform version of the Morris water maze task conducted in 4 blocks of 3 trials over two days shows no significant differences among treatment groups for mice infused with de-oligo or re-oligo tau from sham or shockwave exposed mice (2-way RM ANOVA for latency with treatment and trial as factors shows no significant effect of treatment ($F(3,46) = 1.147$, $P = 0.3403$), but a significant effect of trial ($F(3,138) = 68.56$, $P < 0.0001$. **G)** Swim speed for the indicated groups during the marked platform version of the Morris water maze task shown in (F). 2-way RM ANOVA for speed with treatment and trial as factors shows no significant effects of treatment ($F(3,46) = 0.2953$, $P = 0.8286$), but a significant effect of trial ($F(3,138) = 5.978$, $P = 0.0007$). **H)** Threshold for responses to foot shocks of increasing intensity shows no significant differences among treatment groups for mice infused with de-oligo or re-oligo tau from sham or shockwave exposed mice (2-way RM ANOVA for effect of treatment $F(3,46) = 0.1460$, $P = 0.9317$; Tukey's post-hoc comparisons revealed no significant differences between treatment groups for any response, $P > 0.05$). **I)** Time spent in the center of an open field environment during a 20 minute trial shows no significant differences among groups for mice infused with sham or blast tau that was de-oligomerized by treatment with reducing reagent alone (de-oligo tau) or de-oligomerized then re-oligomerized by peroxide treatment (re-oligo tau) (2-way RM ANOVA for effect of group with group and day as factors: $F(3,46) = 0.2054$, $P = 0.8921$). Moreover, Tukey's multiple comparisons show no significant differences between any groups on either day ($P > 0.05$). N = 13 de-oligo sham tau, 13 de-oligo blast tau, 12 re-oligo sham tau, 12 re-oligo blast tau. **J)** Graph shows no significant difference in distance traveled by the same animals in (I) during a 20-minute exposure to an open field environment (2-way ANOVA for treatment $F(3,46) = 0.2855$, $P = 0.8356$; Tukey's post-hoc pairwise comparisons show no significant differences between groups on either day ($P > 0.05$). All data presented as mean $\pm$ SEM.

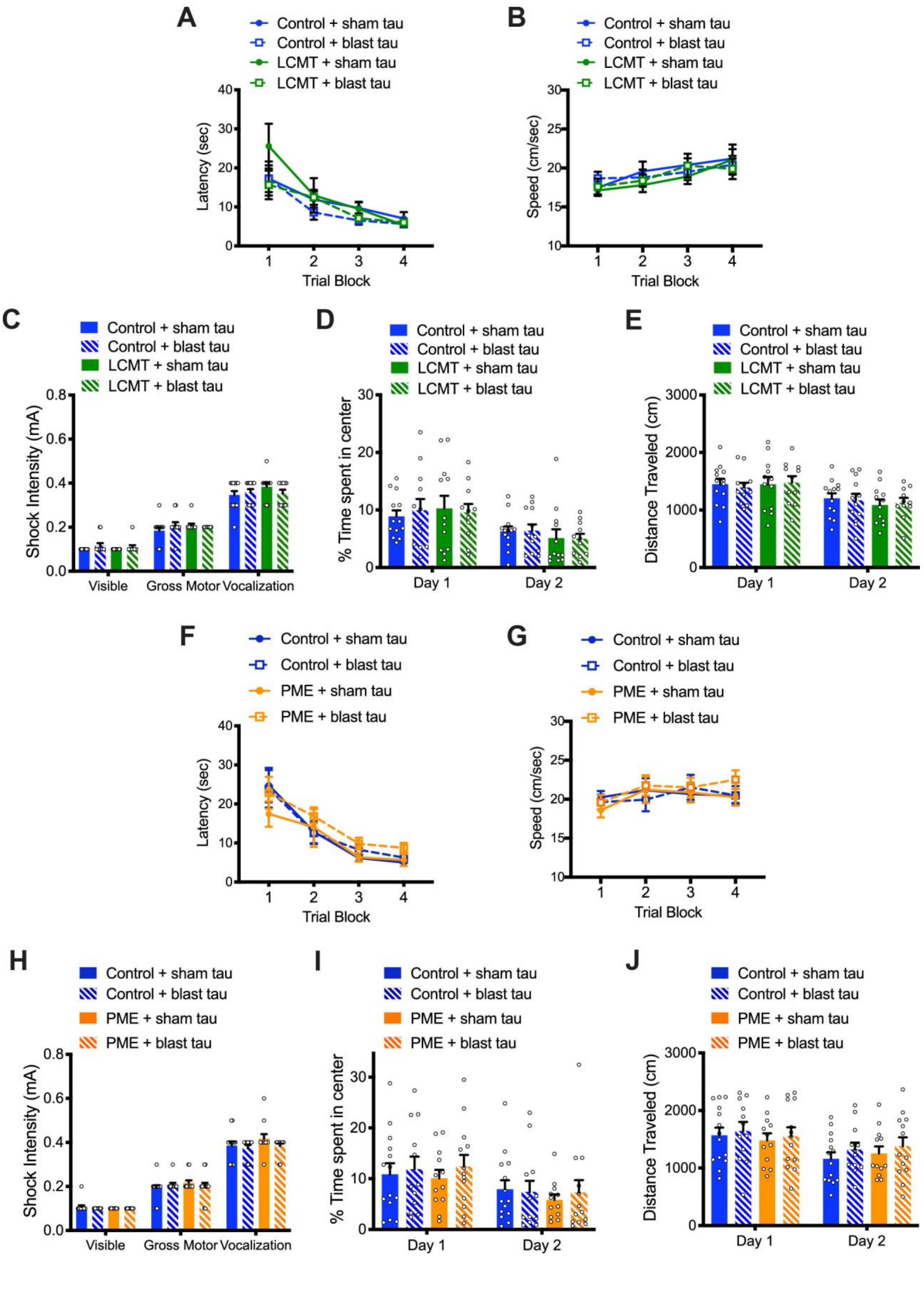
A
Control + sham tau
Control + blast tau
LCMT + sham tau
LCMT + blast tau
Latency (sec)
Trial Block
B
Speed (cm/sec)
Trial Block
C
Shock Intensity (mA)
Visible
Gross Motor
Vocalization
D
% Time spent in center
Day 1
Day 2
E
Distance Traveled (cm)
F
PME + sham tau
PME + blast tau
G
H
I
J

**Fig. S5: Transgenic overexpression of LCMT-1 or PME-1 does not alter the open field behavior, foot shock perception, or marked platform water maze performance of mice treated with tau prepared from shockwave-exposed mice. A)** Time to reach the escape platform in a marked platform version of the Morris water maze task conducted in 4 blocks of 3 trials over two days for LCMT-1 overexpressing transgenic mice and sibling controls infused sham or blast tau shows no significant differences among groups (2-way RM ANOVA for latency with group and trial as factors shows no significant effect of group ($F(3,44) = 0.8984$, $P = 0.4496$), but a significant effect of trial ($F(3,132) = 21.27$, $P < 0.0001$). **B)** Swim speed for the indicated groups during the marked platform version of the Morris water maze task shown in (A). 2-way RM ANOVA for speed with group and trial as factors shows no significant effect of group ($F(3,44) = 0.3871$, $P = 0.7629$), but a significant effect of trial ($F(3,132) = 5.853$, $P = 0.0009$). **C)** Threshold for responses to foot shocks of increasing intensity for LCMT-1 overexpressing transgenic mice and sibling controls infused with sham or blast tau shows no significant differences among groups (2-way RM ANOVA for effect of group $F(3,44) = 1.358$, $P = 0.2680$; Tukey's post-hoc comparisons revealed no significant differences between treatment groups for any response, $P > 0.05$). **D)** Time spent in the center of an open field environment during two 10-minute trials over two days for LCMT-1 overexpressing transgenic mice and sibling controls infused with sham or blast tau shows no significant differences among treatment groups (2-way RM ANOVA for effect of group with group and day as factors: $F(3,44) = 0.0788$, $P = 0.9712$). Moreover, Tukey's multiple comparisons show no significant differences between any groups on either day ($P > 0.05$). N = 13 control+sham tau, 12 control+blast tau, 12 LCMT+sham tau, 11 LCMT+blast tau mice). **E)** Graph shows no significant difference in distance traveled by the same animals in (D) during two 10 minute trials over two days in an open field environment (2-way ANOVA for group with group and day as factors $F(3,44) = 0.0886$, $P = 0.9659$; Tukey's post-hoc pairwise comparisons show no significant differences between groups on either day ($P > 0.05$.). **F)** Time to reach the escape platform in a marked platform version of the Morris water maze task conducted in 4 blocks of 3 trials over two days for PME-1 overexpressing transgenic mice and sibling controls infused with sham or blast tau shows no significant differences among groups (2-way RM ANOVA for latency with group and trial as factors shows no significant effect of group ($F(3,47) = 0.5615$, $P = 0.6431$), but a significant effect of trial ($F(3,141) = 41.69$, $P < 0.0001$). **G)** Swim speed for the indicated groups during the marked platform version of the Morris water maze task shown in (F). 2-way RM ANOVA for speed with group and trial as factors shows no significant effect of group ($F(3,47) = 0.3177$, $P = 0.8125$), and no significant effect of trial ($F(3,141) = 2.357$, $P = 0.0744$). **H)** Threshold for responses to foot shocks of increasing intensity for PME-1 overexpressing transgenic mice and sibling controls infused with sham or blast tau shows no significant differences among groups (2-way RM ANOVA for effect of group $F(3,47) = 1.013$, $P = 0.3954$; Tukey's post-hoc comparisons revealed no significant differences between treatment groups for any response, $P > 0.05$). **I)** Time spent in the center of an open field environment during two 10-minute trials over two days for PME-1 overexpressing transgenic mice and sibling controls infused with sham or blast tau shows no significant differences among treatment groups (2-way RM ANOVA for effect of group with group and day as factors: $F(3,47) = 0.1910$, $P = 0.9020$). Moreover, Tukey's multiple comparisons show no significant differences between any groups on either day ($P > 0.05$). N = 14 control+sham tau, 12 control+blast tau, 12 PME+sham tau, 13 PME+blast tau mice. **J)** Graph shows no significant difference in distance traveled by the same animals in (I) during two 10-minute trials over two days in an open field environment (2-way ANOVA for group with group and day as factors $F(3,47) = 0.2468$, $P = 0.8632$; Tukey's post-hoc pairwise comparisons show no significant differences between groups on either day ($P > 0.05$). All data presented as mean $\pm$ SEM.

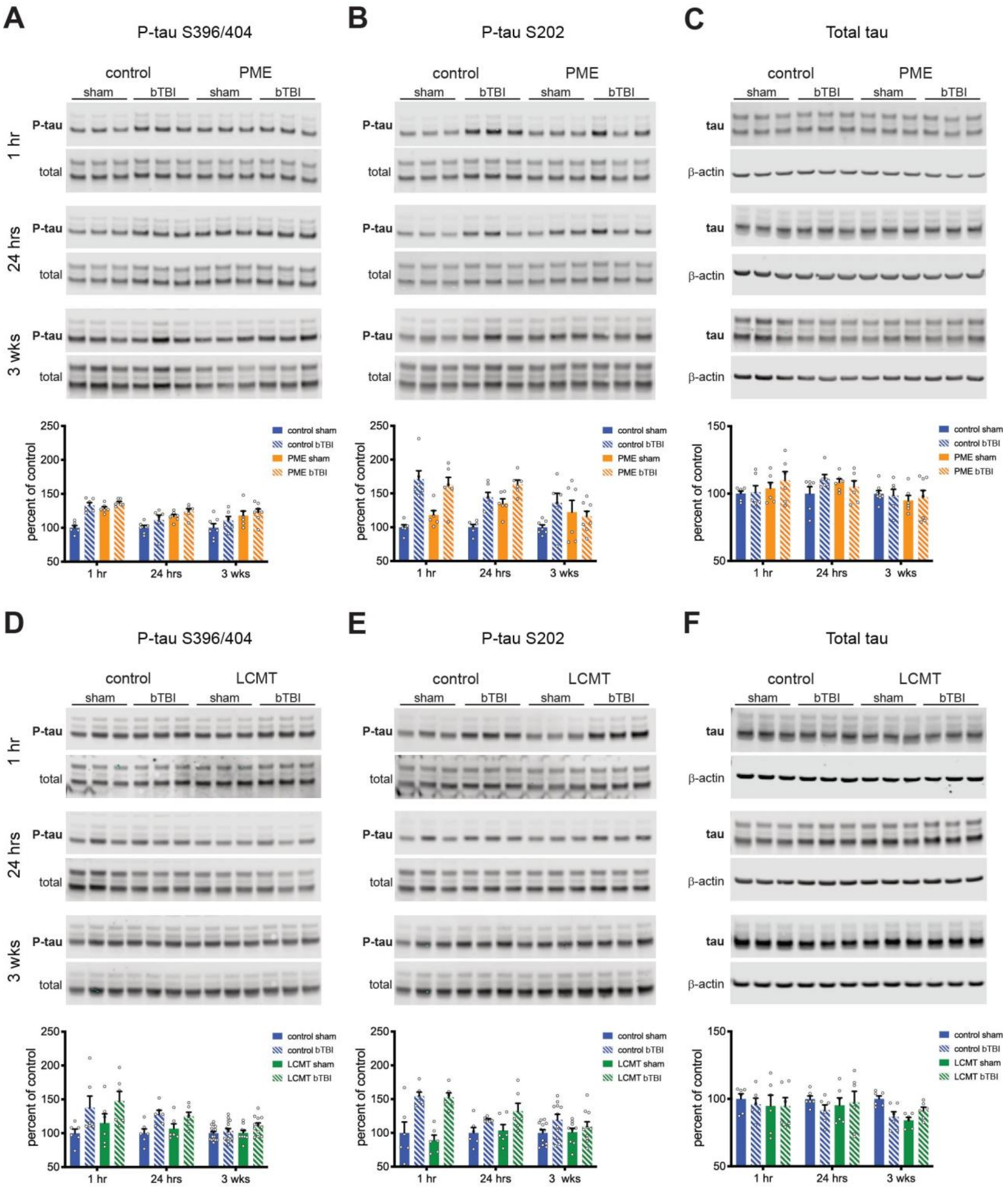


**Fig. S6: Transgenic overexpression of PME or LCMT does not significantly affect tau phosphorylation in shockwave exposed mice.** Representative western blots and quantification of immunoreactivity from hippocampal homogenates from sham and shockwave exposed mice at 1 hr, 24 hrs, and 3 wks after injury. **A)** Immunoreactivity for phosphorylated tau at serine 396/404 and corresponding total tau from sham and shockwave exposed PME-1 overexpressing and control mice. 2-way ANOVA with treatment and genotype as factors at 1 hr shows significant effects of treatment ($F(1,20) = 44.89$, $P < 0.0001$) and genotype ($F(1,20) = 26.91$, $P < 0.0001$), and Dunnett's post-hoc comparisons show significant differences between sham controls and each of the other groups ($P < 0.0001$). 2-way ANOVA with treatment and genotype as factors at 24 hrs shows a significant effect of genotype ($F(1,20) = 9.538$, $P = 0.0058$) but not treatment ($F(1,20) = 9.538$, $P = 0.0654$), and Dunnett's post-hoc comparisons show a significant difference between sham controls and the PME bTBI group ($P = 0.0054$) but neither of the of the other two groups ($P = 0.0614$ for sham PME, 0.2679 for bTBI control). 2-way ANOVA with treatment and genotype as factors at 3 wks shows a significant effect of genotype ($F(1,23) = 6.769$, $P = 0.0159$) but not treatment ($F(1,23) = 1.804$, $P = 0.1924$), and Dunnett's post-hoc comparisons show a significant difference between sham controls and the PME bTBI group ($P = 0.0191$) but neither of the of the other two groups ($P = 0.1129$ for sham PME, 0.4780 for bTBI control). **B)** Immunoreactivity for phosphorylated tau at serine 202 and corresponding total tau from sham and shockwave exposed PME-1 overexpressing and control mice. 2-way ANOVA with treatment and genotype as factors at 1 hr shows a significant effect of treatment ($F(1,20) = 35.23$, $P < 0.0001$) but not genotype ($F(1,20) = 0.1941$, $P = 0.6642$), and Dunnett's post-hoc comparisons show significant differences between sham controls and the control bTBI ($P = 0.0001$) and PME bTBI ($P = 0.0006$) groups but not the PME sham group ($P = 0.4212$). 2-way ANOVA with treatment and genotype as factors at 24 hrs show significant effects of genotype ($F(1,20) = 18.7$, $P = 0.0003$) and treatment ($F(1,20) = 31.99$, $P < 0.0001$), and Dunnett's post-hoc comparisons show significant differences between sham controls and each of the other groups ($P = 0.0024$ vs. sham PME, $P = 0.0003$ vs. control bTBI, $P < 0.0001$ vs. PME bTBI). 2-way ANOVA with treatment and genotype as factors at 3 wks shows no significant effects of genotype ($F(1,23) = 0.01367$, $P = 0.9079$) or treatment ($F(1,23) = 1.720$, $P = 0.2026$), and Dunnett's post-hoc comparisons show no significant differences between sham controls and any of the other groups ($P = 0.3964$ vs. sham PME, $P = 0.0942$ vs. control bTBI, $P = 0.5879$ vs. PME bTBI). **C)** Immunoreactivity for total tau and corresponding $\beta$-actin from sham and shockwave exposed PME-1 overexpressing and control mice. 2-way ANOVA with treatment and genotype as factors at 1 hr shows no significant effects of genotype ($F(1,23) = 1.743$, $P = 0.2016$) or treatment ($F(1,23) = 0.4829$, $P = 0.4951$), and Dunnett's post-hoc comparisons show no significant differences between sham controls and any of the other groups ($P = 0.8960$ vs. sham PME, $P = 0.9986$ vs. control bTBI, $P = 0.3692$ vs. PME bTBI). 2-way ANOVA with treatment and genotype as factors at 24 hrs shows no significant effects of genotype ($F(1,20) = 0.0882$, $P = 0.7696$) and treatment ($F(1,20) = 0.8305$, $P = 0.3730$), and Dunnett's post-hoc comparisons show no significant differences between sham controls and any of the other groups ($P = 0.3570$ vs. sham PME, $P = 0.1766$ vs. control bTBI, $P = 0.7297$ vs. PME bTBI). 2-way ANOVA with treatment and genotype as factors at 3 wks shows no significant effects of genotype ($F(1,23) = 0.5746$, $P = 0.4588$) or treatment ($F(1,23) = 0.01173$, $P = 0.9147$), and Dunnett's post-hoc comparisons show no significant differences between sham controls and any of the other groups ($P = 0.7463$ vs. sham PME, $P = 0.9920$ vs. control bTBI, $P = 0.9296$ vs. PME bTBI). **D)** Immunoreactivity for phosphorylated tau at serine 396/404 and corresponding total tau from sham and shockwave exposed LCMT-1 overexpressing and control mice. 2-way ANOVA with treatment and genotype as factors at 1 hr shows a significant effect of treatment ($F(1,20) = 6.627$, $P = 0.0181$) but not genotype ($F(1,20) = 0.8381$, $P = 0.3709$). However, differences between sham controls and control bTBI ($P = 0.1528$) and LCMT bTBI ($P = 0.0580$) do not reach significance in Dunnett's post-hoc comparisons. 2-way ANOVA with treatment and genotype as factors at 24hrs shows a significant effect of treatment ($F(1,20) = 12.15$, $P = 0.0023$) but not genotype ($F(1,20) = 0.07083$, $P = 0.7929$). Dunnett's post-hoc comparisons show significant differences between sham controls and the control bTBI ($P = 0.0184$) and LCMT bTBI ($P = 0.0396$) groups, but not the LCMT sham group ($P = 0.8049$). 2-way ANOVA with treatment and genotype as factors at 3 wks shows a non-significant trend for effect of treatment (F(1,48) = 3.903, P = 0.0540) but not genotype (F(1,48) = 1.365, P = 0.2484), and Dunnett's post-hoc comparisons show no significant differences between sham controls and any of the other groups (P = 0.9999 vs. LCMT sham, P = 0.8292 vs. control bTBI, P = 0.0628 vs. LCMT bTBI). **E)** Immunoreactivity for phosphorylated tau at serine 202 and corresponding total tau from sham and shockwave exposed LCMT-1 overexpressing and control mice. 2-way ANOVA with treatment and genotype as factors at 1 hr shows a significant effect of treatment (F(1,20) = 34.31, P < 0.0001) but not genotype (F(1,20) = 1.024, P = 0.3236), and Dunnett's post-hoc comparisons show significant differences between sham controls and the control bTBI (P = 0.0029) and LCMT bTBI (P = 0.0038) groups but not the LCMT sham group (P = 0.8029). 2-way ANOVA with treatment and genotype as factors at 24hrs shows a significant effect of treatment (F(1,20) = 8.033, P = 0.0102) but not genotype (F(1,20) = 0.8440, P = 0.3692). Dunnett's post-hoc comparisons a show significant difference between sham controls and the LCMT bTBI group (P = 0.0395) and but not the control bTBI (P = 0.2636) or the LCMT sham (P = 0.9830) groups. 2-way ANOVA with treatment and genotype as factors at 3 wks shows a significant effect of treatment (F(1,38) = 4.180, P = 0.0479) but not genotype (F(1,38) = 0.5061, P = 0.4812), and Dunnett's post-hoc comparisons show no significant differences between sham controls and any of the other groups (P = 0.9991 vs. LCMT sham, P = 0.0951 vs. control bTBI, P = 0.6622 vs. LCMT bTBI). **F)** Immunoreactivity for total tau and corresponding $\beta$-actin from sham and shockwave exposed LCMT-1 overexpressing and control mice. 2-way ANOVA with treatment and genotype as factors at 1 hr shows no significant effects of treatment (F(1,20) = 0.1306, P = 0.7216) or, genotype (F(1,20) = 0.3136, P = 0.5817), and Dunnett's post-hoc comparisons show no significant differences between sham controls and any of the other groups (P = 0.8678 vs. sham LCMT, P = 0.9309 vs. control bTBI, P = 0.8525 vs. LCMT bTBI). 2-way ANOVA with treatment and genotype as factors at 24 hrs shows no significant effects of genotype (F(1,20) = 0.008026, P = 0.9295) or treatment (F(1,20) = 0.3219, P = 0.5768), and Dunnett's post-hoc comparisons show no significant differences between sham controls and any of the other groups (P = 0.8835 vs. sham LCMT, P = 0.5953 vs. control bTBI, P = 0.9740 vs. LCMT bTBI). 2-way ANOVA with treatment and genotype as factors at 3 wks shows no significant effects of genotype (F(1,20) = 3.955, P = 0.0606) or treatment (F(1,20) = 1.233, P = 0.2799), and Dunnett's post-hoc comparisons show no significant differences between sham controls and any of the other groups (P = 0.5642 vs. sham LCMT, P = 0.8268 vs. control bTBI, P > 0.9999 vs. LCMT bTBI). All data presented as mean $\pm$ SEM.

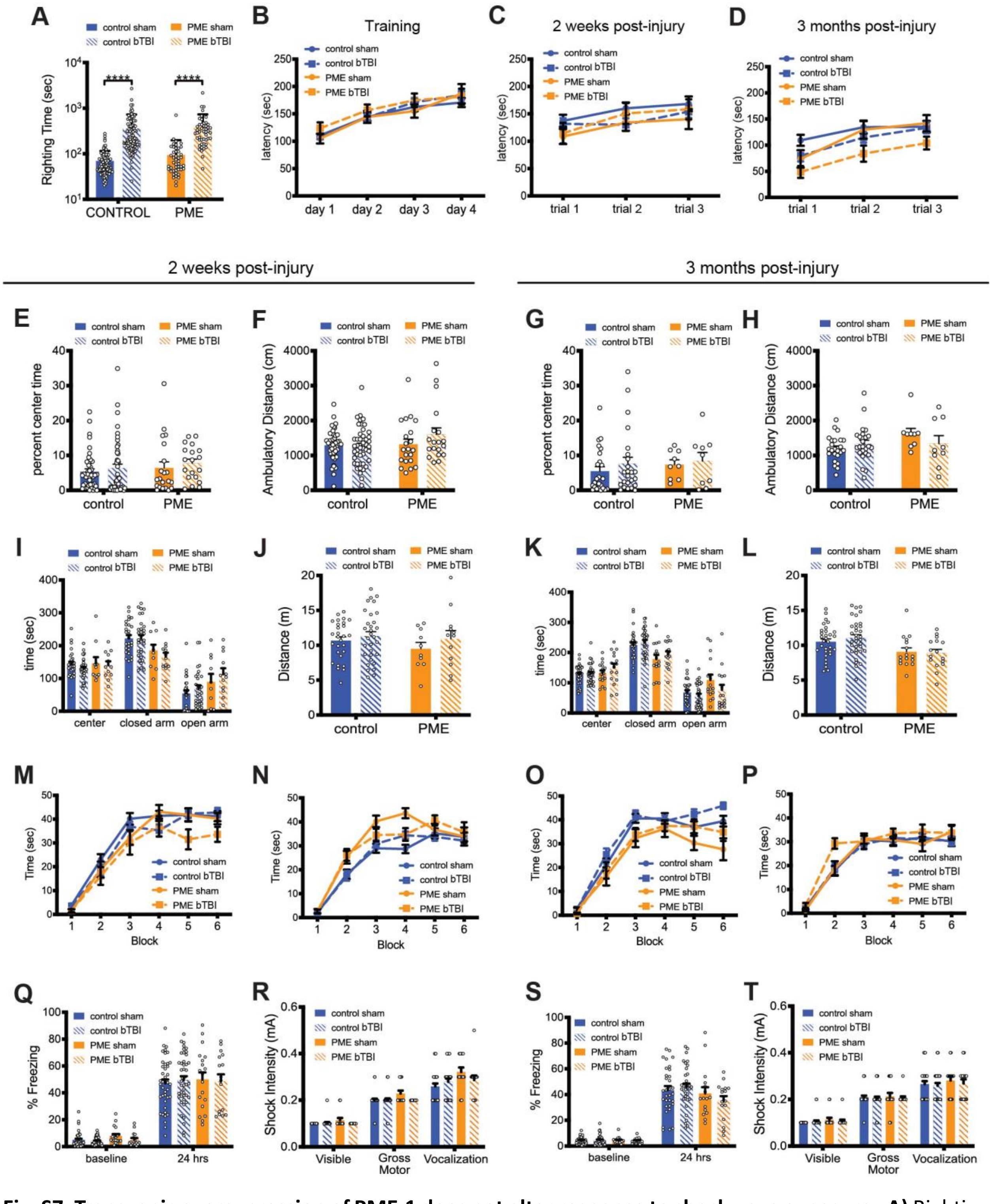


**Fig. S7: Transgenic overexpression of PME-1 does not alter response to shockwave exposure. A)** Righting time for shockwave (bTBI) and sham exposed PME transgenic and control mice showed a significant effect of treatment (2-way ANOVA F(1,184) = 62.04, P < 0.0001, N = 47 control sham, 47 control bTBI, 46 PME sham, 48 PME bTBI; Tukey's comparisons of sham vs. bTBI controls, and sham vs. bTBI PME animals: P < 0.0001), but not genotype (2-way ANOVA F(2,184) = 1.123, P = 0.2907; Tukey's comparisons of: sham controls vs. sham PME, P = 0.9571; and bTBI controls vs. bTBI PME, P = 0.7535). **B)** Latency to fall on an accelerating rotarod task for PME overexpressing mice and controls during two training trials on each of 4 days prior to shockwave exposure or sham treatment shows no significant difference between groups (2-way RM ANOVA $F(3,225) = 0.4128$, $P = 0.7440$), but a significant effect of training day ($F(3,675) = 105.3$, $P < 0.0001$). N = 76 control sham, 81 control bTBI, 37 PME sham, 35 PME bTBI. **C)** Latency to fall on an accelerating rotarod task during each of three training trials performed 13 days after shockwave or sham exposure shows no significant difference between groups (2-way ANOVA for effect of group $F(3,124) = 0.8675$, $P = 0.4600$; trial $F(2,248) = 16.73$, $P < 0.0001$; N = 43 control sham, 46 control bTBI, 21 PME sham, 18 PME bTBI). **D)** Latency to fall on an accelerating rotarod task during each of three training trials performed 3 months after shockwave or sham exposure shows no significant differences among groups (2-way RM ANOVA for effect of group $F(3,94) = 2.598$, $P = 0.0568$; trial $F(2,188) = 61.76$, $P < 0.0001$; N = 32 control sham, 34 control bTBI, 16 PME sham, 16 PME bTBI). **E)** Graph shows no significant differences in time spent in the center of an open field environment during a 20 minute trial for shockwave or sham exposed PME overexpressing mice and controls 13 days after injury (2-way ANOVA for effect of treatment $F(1,125) = 0.8500$, $P=0.3583$; genotype $F(1,125) = 0.9478$, $P = 0.3322$, interaction $F(1,125) = 0.01532$, $P = 0.9017$; Tukey's post-hoc pairwise comparisons show no significant differences between groups $P < 0.05$; N = 44 control sham, 45 control bTBI, 21 PME sham, 19 PME bTBI). **F)** Graph shows no significant difference in distance traveled by the same animals in (E) during a 20 minute exposure to an open field environment (2-way ANOVA for effect of treatment $F(1,125) = 1.324$, $P = 0.2520$; genotype $F(1,125) = 2.645$, $P = 0.1064$, interaction $F(1,125) = 1.803$, $P = 0.1818$). **G)** Graph shows no significant differences in time spent in the center of an open field environment during a 20 minute trial for sham or shockwave exposed PME overexpressing mice and controls 3 months after injury (2-way ANOVA for effect of treatment $F(1,62) = 0.5607$, $P = 0.4568$; genotype $F(1,62) = 0.4204$, $P = 0.5191$, interaction $F(1,62) = 0.07459$, $P = 0.7857$; N = 24 control sham, 24 control bTBI, 9 PME sham, 9 PME bTBI). **H)** Graph shows no significant differences in distance traveled by the same animals in (G) during a 20 minute exposure to an open field environment (2-way ANOVA for effect of treatment $F(1,62) = 0.2739$, $P = 0.6026$; genotype $F(1,62) = 2.461$, $P = 0.1218$, interaction $F(1,62) = 2.345$, $P = 0.1307$). **I)** Graph shows time spent in the open arms, closed arms, and center of an elevated plus maze for shockwave and sham exposed PME overexpressing mice and controls 14 days after injury. 2-way ANOVA shows no significant effect of treatment ($F(1,77) = 1.831$, $P = 0.1800$), but a significant effect of genotype ($F(1,77) = 7.553$, $P = 0.0075$) suggesting modestly reduced anxiety in the PME overexpressing animals. However, 2-way ANOVA showed no significant interaction ($F(1,77) = 0.0988$, $P = 0.7541$), and Tukey's pairwise comparisons showed a significant difference only between the sham controls and PME bTBI groups ($P = 0.0198$). N = 28 control sham, 31 control bTBI, 10 PME sham, 12 PME bTBI. **J)** Graph shows no significant difference in distance traveled by the same animals in (I) during elevated plus maze testing (2-way ANOVA for effect of treatment $F(1,77) = 1.482$, $P = 0.2272$; genotype $F(1,77) = 0.910$, $P = 0.3455$, interaction $F(1,77) = 0.2181$, $P = 0.6418$; Tukey's post-hoc pairwise comparisons show no significant differences between genotype/treatment groups, $P > 0.05$). **K)** Graph shows time spent in the open arms, closed arms, and center of an elevated plus maze for shockwave and sham exposed PME overexpressing mice and controls 3 months after injury. 2-way ANOVA shows no significant effect of treatment ($F(1,93) = 3.918$, $P = 0.0507$), but a significant effect of genotype ($F(1,93) = 7.437$, $P = 0.0076$) suggesting modestly reduced anxiety in the PME overexpressing animals. 2-way ANOVA showed no significant interaction ($F(1,93) = 0.9537$, $P = 0.3313$), and Tukey's pairwise comparisons showed a significant difference between only the control bTBI and PME sham groups ($P = 0.0062$). N = 31 control sham, 34 control bTBI, 16 PME sham, 16 PME bTBI). **L)** Graph shows distance traveled by the same animals in (K) during elevated plus maze testing. 2-way ANOVA shows no significant effect of treatment (F(1,93) = 0.1145, P = 0.7358), or interaction (F(1,93) = 0.2575, P = 0.7734), but a significant effect of genotype (F(1, 93) = 12.30, P < 0. 0007). **M)** Time immobile during 1 min blocks on a forced swim task for PME over expressing mice and controls 14 days after shockwave or sham exposure. 2-way RM ANOVA shows a significant effect of time (F(5,450) = 174.6, P < 0.0001) and also group (F(3,90) = 2.739, P = 0.0480), but Tukey's pairwise comparisons suggest a significant difference between only the control sham and PME bTBI groups (P = 0.0263). N = 32 control sham, 33 control bTBI, 14 PME sham, 15 PME bTBI. **N)** Time immobile during 1 min blocks on a tail suspension task for PME overexpressing mice and controls 17 days after shockwave or sham exposure. 2-way RM ANOVA shows significant effect of time (F(5,450) = 208.6, P < 0.0001), and group (F(3,90) = 4.588, P = 0.0049), but Tukey's pairwise comparisons suggest significant differences between only the control sham and PME sham (P = 0.0155) and control bTBI and PME sham (P = 0.0347) groups. N = 31 control sham, 34 control bTBI, 14 PME sham, 15 PME bTBI. **O)** Time immobile during 1 min blocks on a forced swim task for PME overexpressing mice and controls 3 months after shockwave or sham exposure. 2-way RM ANOVA shows a significant effect of time (F(5,470) = 175.0, P < 0.0001) and also group (F(3,94) = 5.751, P = 0.0012), but Tukey's pairwise comparisons suggest significant differences between only the control sham and PME sham (P = 0.0459) and control bTBI and PME sham (P = 0.0011) groups. N = 32 control sham, 34 control bTBI, 16 PME sham, 16 PME bTBI. **P)** Time immobile during 1 min blocks on a tail suspension task for PME overexpressing mice and controls 3 months after shockwave or sham exposure. 2-way RM ANOVA shows significant effect of time (F(5,440) = 137.1, P < 0.0001), but not group (F(3,88) = 1.945, P = 0.1282). N = 29 control sham, 31 control bTBI, 16 PME sham, 16 PME bTBI. **Q)** Graph of percent of time spent freezing during initial exposure to the training context (baseline) and 24 hrs after foot shock for PME-1 overexpressing transgenic mice and sibling controls 15 days after shockwave or sham exposure. ANOVA for freezing showed no significant differences among groups at 24 hrs (F(3,116) = 0.1846, P = 0.9067), but a significant difference at baseline (F(3,116) = 2.813, P = 0.0424). However, Tukey's pairwise comparisons of baseline freezing responses show a significant difference only between control bTBI and PME sham groups. N = 41 control sham, 44 control bTBI, 19 PME sham, 16 PME bTBI. **R)** Graph of threshold for responses to foot shocks of increasing intensity for PME-1 overexpressing transgenic mice and sibling controls 20 days after shockwave or sham exposure revealed no significant differences in foot-shock perception (one-way ANOVA for first visible response: F(3,91) = 2.490, P = 0.0653; for first gross motor response: F(3,91) = 2.299, P = 0.0827, for first vocal response: F(3,91) = 2.354, P = 0.0772). N = 32 control sham, 34 control bTBI, 14 PME sham, 15 PME bTBI. **S)** Graph of percent of time spent freezing during initial exposure to the training context (baseline) and 24 hrs after foot shock for PME-1 overexpressing transgenic mice and sibling controls 3 months after shockwave or sham exposure. ANOVA for freezing showed no significant differences among groups at 24 hrs (F(3,94) = 1.460, P = 0.2305), or baseline (F(3,94) = 0.09039, P = 0.9652). N = 32 control sham, 34 control bTBI, 16 PME sham, 16 PME bTBI. **T)** Graph of threshold for responses to foot shocks of increasing intensity for PME-1 overexpressing transgenic mice and sibling controls 3 months after shockwave or sham exposure revealed no significant differences in foot-shock perception (one-way ANOVA for first visible response: F(3,94) = 1.200, P = 0.3141; for first gross motor response: F(3, 94) = 0.1894, P = 0.9034, for first vocal response: F(3,94) = 0.4040, P = 0.7504). N = 32 control sham, 34 control bTBI, 16 PME sham, 16 PME bTBI. All data presented as mean ± SEM.

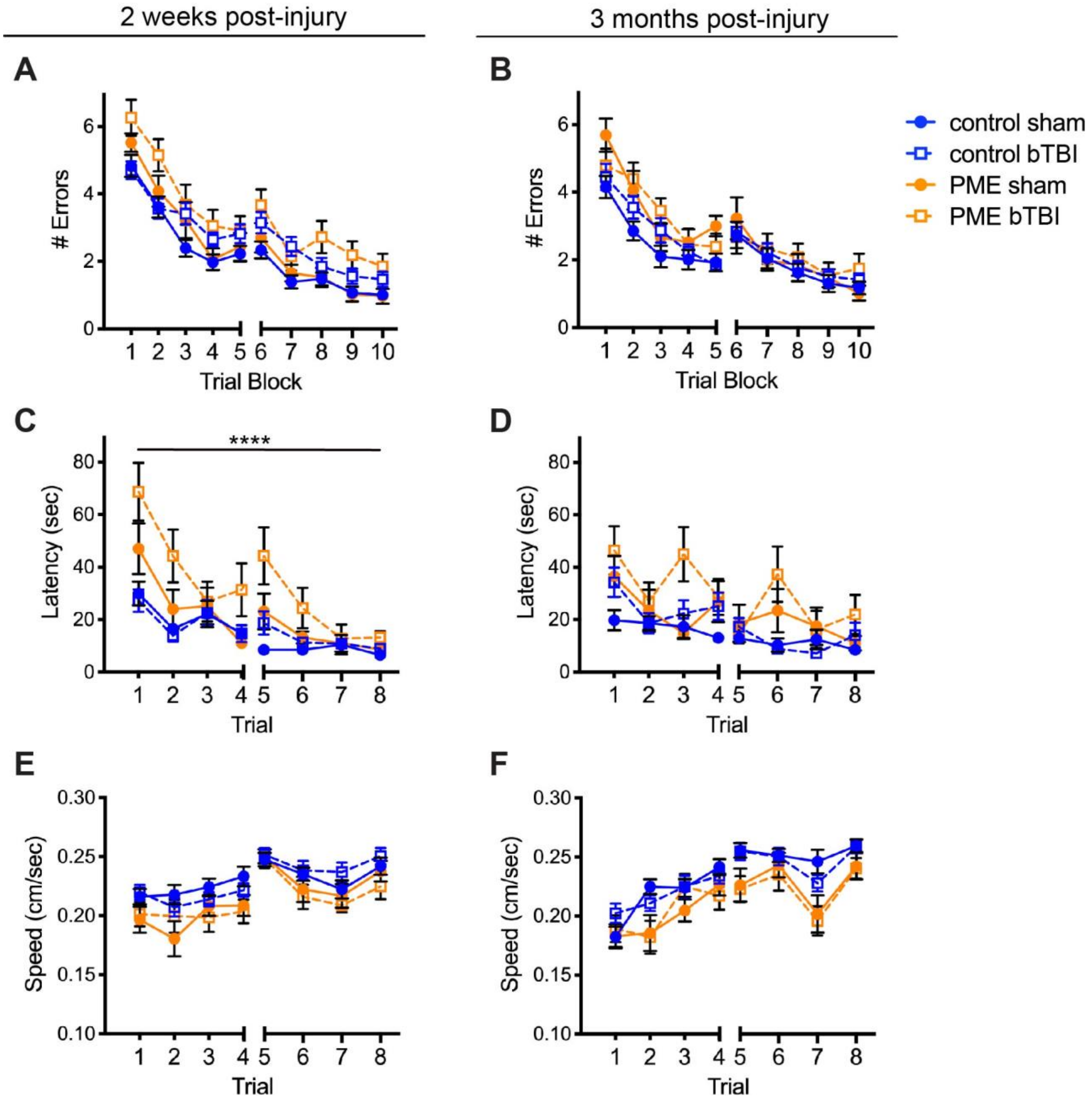


**Fig. S8: Transgenic overexpression of PME does not significantly affect the performance of shockwave exposed mice in a two-day RAWM, but may impair performance of shockwave exposed mice on marked platform Morris water maze task. A)** Number of errors committed during each 3-trial training block of a 2-day RAWM task for PME-1 overexpressing transgenic mice and sibling controls 16 days after shockwave (bTBI) or sham exposure. 2-way RM ANOVA for errors on day 2 (blocks 6-10) with group and block as factors shows no significant differences among groups ($F(3,124) = 3.825$, $P = 0.0116$). N = 43 control sham, 46 control bTBI, 21 PME sham, 18 PME bTBI. **B)** Number of errors committed during each 3-trial training block of a 2-day RAWM task for PME-1 overexpressing transgenic mice and sibling controls 3 months after shockwave or sham exposure. 2-way RM ANOVA for errors on day 2 with group and block as factors shows no significant differences among groups ($F(3,94) = 0.2313$, $P = 0.2313$). N = 32 control sham, 34 control bTBI, 16 PME sham, 16 PME bTBI. **C)** Time to reach the escape platform for PME-1 overexpressing transgenic mice and sibling controls 22 days after shockwave or sham exposure during each of 8 trials conducted over two successive days in a marked platform version of the Morris water maze task. 2-way RM ANOVA for latency with group and trial as factors shows significant effects of group ($F(3,91) = 13.27$, $P < 0.0001$), trial ($F(7,637) = 22.35$, $P < 0.0001$). Dunnett's multiple comparisons show that the PME bTBI group was significantly different from each of the other groups (vs. control sham, $P < 0.0001$; vs. control bTBI, $P < 0.0001$; vs. PME sham, $P = 0.0022$). N = 32 control sham, 34 control bTBI, 14 PME sham, 15 PME bTBI. **D)** Time to reach the escape platform for PME-1 overexpressing transgenic mice and sibling controls 3 months after shockwave or sham exposure during each of 8 trials conducted over two successive days in a marked platform version of the Morris water maze task. 2-way RM ANOVA for latency with group and trial as factors also shows a significant effect of group ($F(3,94) = 7.948$, $P < 0.0001$). Dunnett's multiple comparisons show that the PME bTBI group was significantly different from the control sham ($P < 0.0001$) and control bTBI ($P = 0.0021$) groups, but not the PME sham group ($P = 0.0792$). N = 32 control sham, 34 control bTBI, 16 PME sham, 16 PME bTBI. **E)** Swim speed for animals shown in (C) during each of 8 trials conducted over two successive days in a marked platform version of the Morris water maze task suggests a small but significant reduction in swim speed in PME overexpressing mice. 2-way RM ANOVA for speed with group and trial as factors shows significant effects of group ($F(3,91) = 3.506$, $P = 0.0185$). Dunnett's multiple comparisons show that the PME bTBI group was significantly different from the control sham ($P = 0.0391$) and control bTBI ($P = 0.0346$) groups, but not the PME sham group ($P = 0.9759$). **F)** Swim speed for animals shown in (D) during each of 8 trials conducted over two successive days in a marked platform version of the Morris water maze task also suggests a small but significant reduction in swim speed in PME overexpressing mice. 2-way RM ANOVA for speed with group and trial as factors shows a significant effect of group ($F(3,94) = 4.354$, $P = 0.0064$). Dunnett's multiple comparisons show that the PME bTBI group was significantly different from the control sham ($P = 0.0194$) and control bTBI ($P = 0.0417$) groups, but not the PME sham group ($P = 0.9997$). All data presented as mean $\pm$ SEM.
